# Dynamical memory underlies prolonged plasmid persistence after transient antibiotic treatment

**DOI:** 10.1101/2025.10.31.685803

**Authors:** Zhengqing Zhou, Andrea Weiss, Zhixiang Yao, Xiaoli Chen, Kristen Lok, Hye-in Son, Lingchong You

## Abstract

Plasmids play critical roles in spreading and maintaining antimicrobial resistance (AMR). They often exhibit prolonged persistence upon antibiotic treatment, even when they impose substantial burden on their hosts. This persistence has been primarily attributed to rapid horizontal transfer or low plasmid cost. However, these mechanisms cannot account for the slow decay of burdensome plasmids with poor mobility. Here, we show that the decoupling of time scales between slow segregation loss and fast growth competition leads to a slow-down in plasmid abundance decay at high initial plasmid abundance, reminiscent of the *ghost effect* from nonlinear dynamical systems. Integrating theory, simulations, and quantitative experiments across clonal populations and multi-species bacterial communities, we demonstrate that a transient antibiotic pulse can eliminate plasmid-free cells and create a *ghost* state that extends plasmid persistence from days to months. Our research reveals a generalizable mechanism for the prolonged ecological memory of antibiotic exposure and underscores the need for proactive strategies to curb the spread of AMR.

## Introduction

Plasmids are self-replicating genetic elements that play a critical role in bacterial evolution by enabling the acquisition, maintenance, and dissemination of adaptive traits such as antimicrobial resistance (AMR)^1–3^, giving rise to many multidrug-resistant high-risk clones^4,5^ and accelerating the global AMR crisis^3^. Although plasmid carriage is often assumed to be burdensome in the absence of selection, leading to rapid plasmid loss^6,7^, empirical studies suggest otherwise. Resistance frequency often remains high for months after short-term antibiotic treatments (days to weeks), despite the absence of continued selective pressure thereafter^8–12^. Given that many resistance determinants reside on plasmids^11,13–19^, the slow resistance reversal suggests burdensome plasmid can still experience long transients after antibiotic exposure.

Different mechanisms may account for this prolonged persistence after antibiotic treatment. For instance, positive selection can increase the frequency of compensatory mutations that reduce the cost of plasmid carriage and slow down the decay of plasmids^20^. Long half-life of certain antibiotics can exert sustained selection for resistance after the treatment ceased^9^. More broadly, the persistence or slow reversal of resistance or plasmid carriage can be associated with low or zero fitness cost^6,21–23^, high conjugation rate^24–26^, or both^24,27^. Co-selection by antibiotics^6,28^ or environmental contaminants^29^ on the same mobilizable gene element has also been proposed. However, these mechanisms cannot account for the slow decay of plasmids whose baseline abundance is low, i.e., burdensome, low-mobility plasmids get lost fast.

Past theories suggest another under-appreciated possibility: the slow segregation loss of plasmids^30,31^. While largely ignored due to its relatively minor contribution to plasmid steady state persistence^24,25,27^, slow segregation loss can sustain plasmid carriage in clones for a long time ^32,33^, before plasmid-encoded high-burden cargoes drive the plasmid to get completely lost. This provides a theoretical basis for the prolonged plasmid carriage following antibiotic exposure: antibiotics can eliminate the plasmid-free competitor cells, so the plasmid needs to go through the slow segregation loss process before rapidly decrease in their abundance due to the growth competition from plasmid-free segregant.

Here, we combine theory, numerical simulations, and quantitative experiments to demonstrate this principle in clonal populations, synthetic *E. coli* communities, and multi-species communities of hospital sink isolates: the decoupling of time scales between segregation loss and growth competition can lead to the observed prolonged ecological memory of plasmids. This principle underscores the need to develop active intervention strategies beyond standard antibiotic stewardship.

## Results

### Population ghost effect slows down plasmid abundance decay

We start from a simple ordinary differential equation (ODE) model depicting the dynamics of a non-mobilizable plasmid within a single strain population (**Methods**). The dynamics of plasmid loss in a population are mainly dictated by two processes: *(1) segregation loss*, where a daughter cell fails to inherit the plasmid during cell division (**Fig. 1a**, pink), and *(2) growth competition*, where plasmid-carrying cells are outcompeted by plasmid-free population due to plasmid-imposed metabolic burden on their host (**Fig. 1a**, purple). Segregation loss is generally a slow process, where different plasmids can deploy active partitioning, post-segregational killing, and high copy-number to minimize its loss during cell division^34,35^. Meanwhile, growth competition operates on a faster timescale.

**Figure 1.**
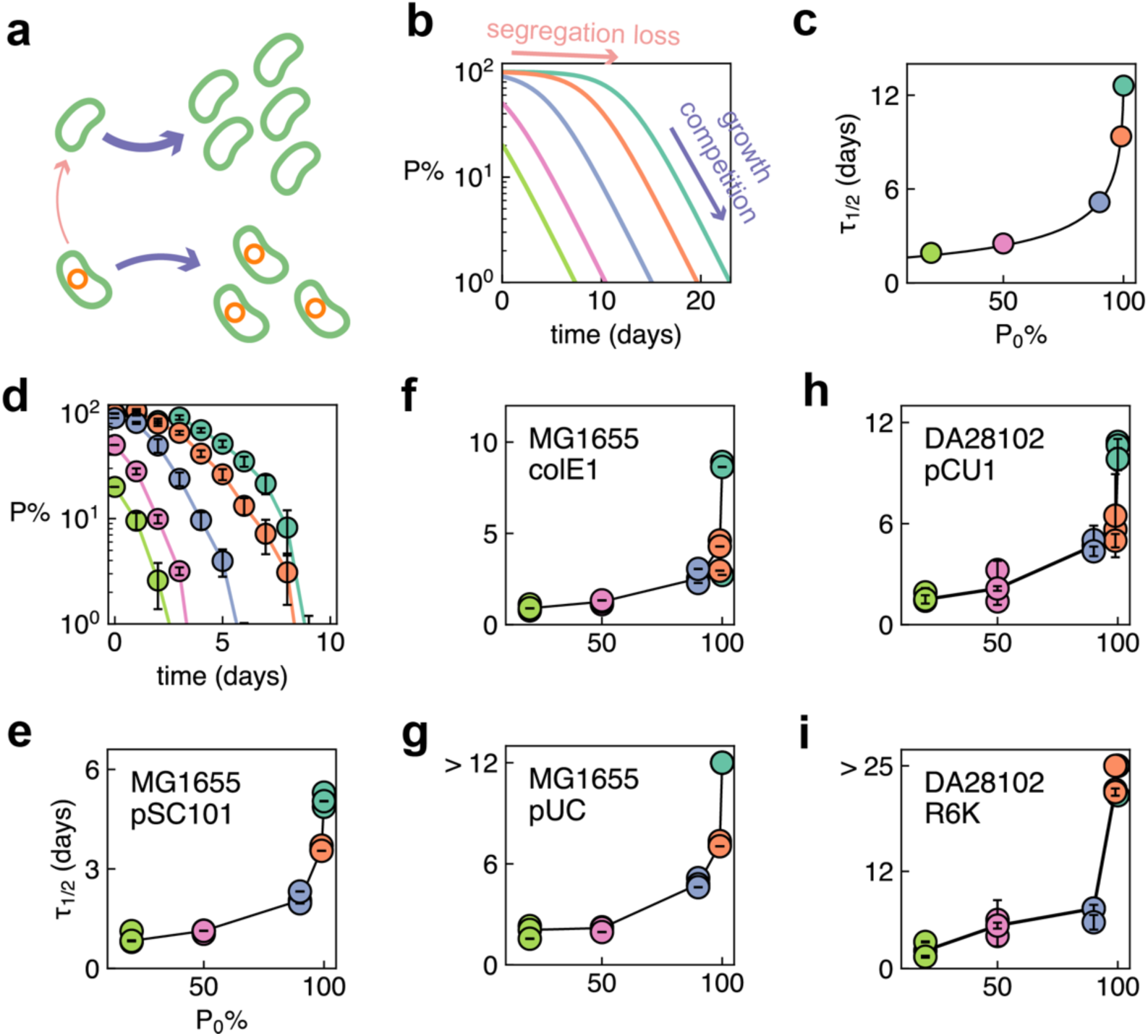
High plasmid abundance nonlinearly increases plasmid half-lives. **(a).** Plasmid abundance decreases through two processes operating on different time scales, slow segregation loss of the plasmid (pink) and rapid out-competition of plasmid-carrying cells by plasmid-free cells (purple). **(b).** Simulated time series of plasmid abundance decay. When starting from high initial abundance P_0_% (teal: 100%, orange: 99%), plasmid abundance exhibits biphasic decay, with the slow phase driven by segregation loss (pink arrow) followed a faster phase driven by growth competition (purple arrow) later. When starting from lower P_0_% values (dusty blue: 90%, pink: 50%, and lime green: 20%), the plasmid abundance follows exponential decay. **(c).** Dependence of plasmid half-life τ_1/2_ on the initial abundance P_0_%. The scatter points represent simulation results; the curve shows the best fit to Eq. (1) in **Methods**. **(d).** Time series of average pSC101 plasmid abundance decay (mean ± SE, n = 3) in *E. coli* MG1655 across different P_0_% (teal: 100%, orange: 99%, dusty blue: 90%, pink: 50%, and lime green: 20%). **(e - g).** Dependence of plasmid half-life τ_1/2_ (mean ± SE, delta method) on the initial abundance P_0_% for non-mobilizable plasmids **(e)** pSC101, **(f)** colE1, and **(g)** pUC. Curves connect the median half-lives at each given initial abundance P_0_%. Right-censored datapoints (τ_1/2_ > 12 days) are shown without error bar. **(h & i).** Dependence of plasmid half-life τ_1/2_ (mean ± SE, delta method) on the initial abundance P_0_% for conjugative plasmids **(h)** pCU1 and **(i)** R6K. Curves connect the median half-lives at each initial abundance P_0_%. Right-censored datapoints (τ_1/2_ > 25 days) are shown without error bar.

We used the ODE model to simulate plasmid abundance (fraction of plasmid carrying cells in a population) over time (**Fig. 1b**), across different initial plasmid abundances of P_0_% = 100%, 99%, 90%, 50%, and 20%. When starting from a pure plasmid-carrying population (P_0_% = 100%), plasmid abundance decays in two phases, corresponding to the two processes: an initial slow decay driven by segregation loss, followed by a faster decay driven by growth competition once plasmid-free cells have emerged. This two-phase behavior is consistent with previous experimental observations^32,36^, where the presence of post-segregational killing mechanism results in a long plateau in which plasmid abundance remains ∼100%. In the absence of active partitioning or post-segregational killing mechanisms, this plateau was significantly shortened or even eliminated. As the initial plasmid abundance decreases, growth competition becomes more prominent, driving the plasmid abundance decay exponentially. Based on a previous work^30^, we can analytically derive the half-life of plasmid abundance decay (**Methods**) which nonlinearly increases with initial plasmid abundance (**Fig. 1c**). When P_0_% << 100%, the abundance decays exponentially with half-life 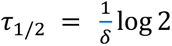. When P_0_% = 100%, plasmid half-life becomes 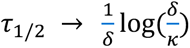, where δ is the rate of growth competition between the plasmid-free and plasmid-carrying populations, and κ is the segregation loss rate. In most cases, κ ≪ δ ^24^, thus half-life τ_1/2_ becomes large, and the high abundance state appear as the *ghost* of an equilibrium^37^. This logarithmic scaling between κ and τ_1/2_ resembles the *ghost effect* coined by Strogatz^37^, originally referring to the loss of stability in a dynamical system during saddle-node bifurcation. The prolonged plasmid carriage in our case instead arises from the remnant of a saddle point (when κ increases from 0 to a small number).

To test our prediction of diverging half-lives near pure population, we monitored the abundance of three non-mobilizable plasmids with different origin of replication, pSC101 (kanamycin resistant, Kan^R^), colE1 (Kan^R^), and pUC (spectinomycin, Spec^R^), all encoding sfGFP, in *E. coli* strain MG1655 over time (**Fig. 1d & Fig. S1**). To facilitate high-throughput measurements, we used the GFP/OD ratio to quantify plasmid abundance (calibration details in **Methods**). Plasmid-carrying MG1655 cells were mixed with plasmid-free MG1655 cells at different volume ratios to generate populations where plasmid carrying cells make up of 100%, 99%, 90%, 50% or 20% of the total population. Populations were cultured in LB and diluted 1:500 every 24 hours (**Methods**). As predicted by the mathematical model, a plateau in plasmid abundance was observed at high initial abundances before exponential decay (**Fig. S1b**), significantly prolonging the half-life of plasmid abundance. Thus, the initial purity of the community, in terms of the plasmid-carrying cell population, dictates the plasmid half-life (**Fig. 1 e - g**). Interestingly, at P_0_% = 100%, pUC persisted in the community. As a high copy plasmid^38^, the stochastic chance of pUC to have segregation error becomes extremely small, preventing the birth of plasmid-free cells and the subsequent growth competition. Moreover, for the populations with plasmid abundance decay (pUC with P_0_% = 99% and 90%, and colE1 with P_0_% = 99%), the plasmid abundances sometimes rebounded, suggesting potential compensatory mutations.

We further quantified the dynamics of two conjugative plasmids, pCU1 and R6K in *E. coli* DA28102, with daily 100-fold dilution in LB media supplemented with 25 μg/mL chloramphenicol (Cm) to prevent contamination. Under these conditions, pCU1 and R6K cannot persist at steady state and were lost rapidly for P_0_% < 90%. With P_0_% = 90%, pCU1 had a half-life of 4.7 ± 0.4 days (mean ± SD, n = 3), and R6K had a half-life of 6.8 ± 1.0 days (mean ± SD, n = 3). However, both exhibited drastically prolonged persistence when P_0_% is near 100% (**Fig. 1h & i, Fig. S2**).

### Transient antibiotic selection extends plasmid half-life through purifying selection

Antibiotics can eliminate plasmid-free cells in a population and purify the population to enter the ghost effect region (**Fig. 2a**). We thus hypothesize selection with antibiotics could prolong plasmid persistence. We assembled populations of *E. coli* MG1655 with an equal mixture of plasmid-free and plasmid-carrying cells (P_0_%=50%). We used the non-mobilizable, sfGFP-encoding plasmids pSC101, colE1, and pUC for high-throughput measurement. We applied a pulse of antibiotic selection to the populations between day 2 and 3. After growing the population in LB supplemented with a gradient of antibiotics (Kan for pSC101 and colE1, and Spec for pUC) for 24 hours, we removed the selection and tracked the GFP/OD readout over time (**Fig. 2b – d, Fig. S3b**). The half-lives of the three plasmids all substantially increased with the strength of the pulsed antibiotic selection (**Fig. 2e - g**), and plateaued above critical concentrations. The critical concentrations are 3.125 µg/mL Kan for pSC101, 6.25 µg/mL Kan for colE1, and 12.5 µg/mL Spec for pUC.

**Figure 2.**
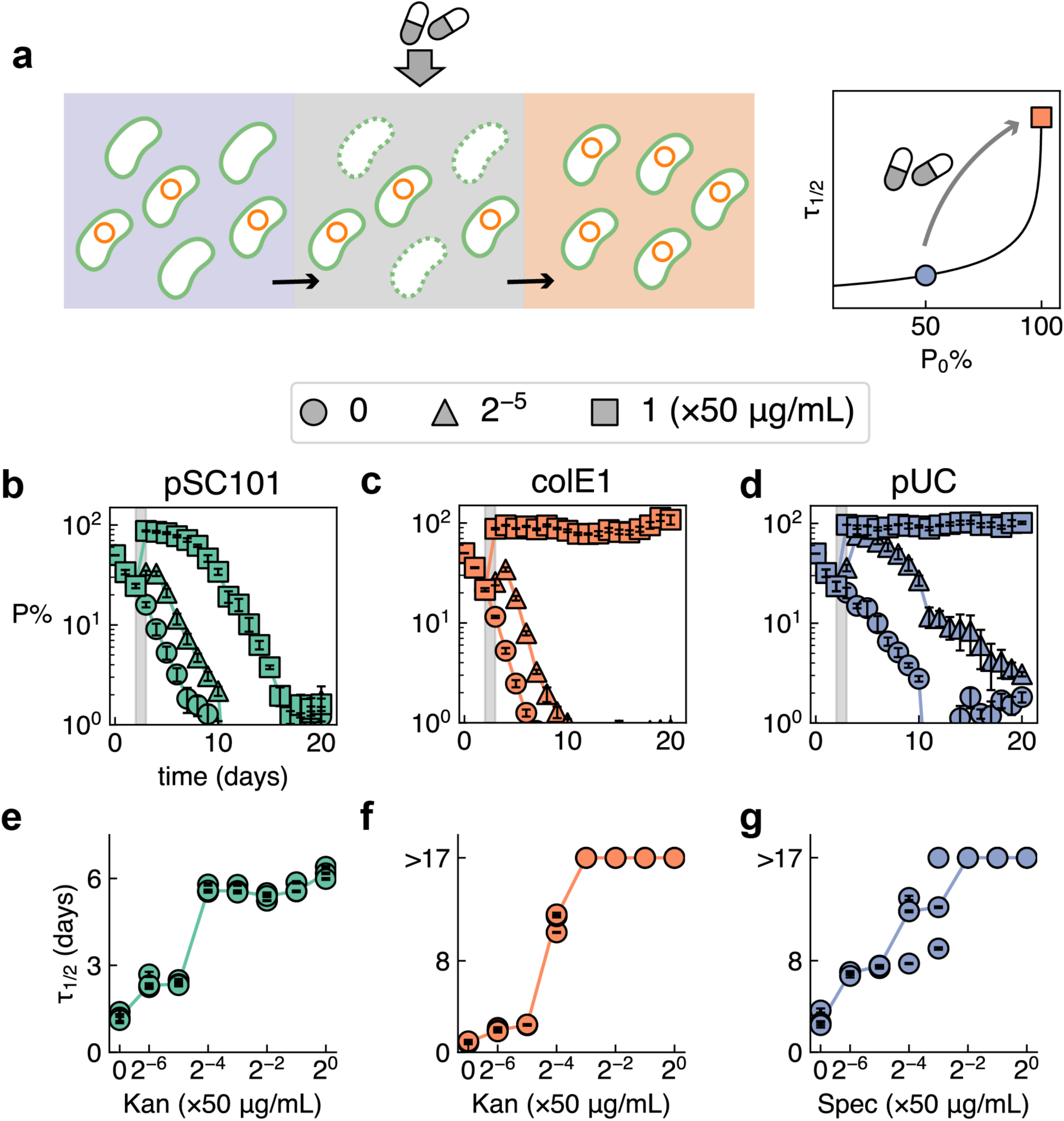
A transient antibiotic pulse prolongs plasmid persistence by inducing ghost effect. **(a).** In a mixed population containing both plasmid-carrying and plasmid-free cells (purple), an antibiotic purifies the population by eliminating the plasmid-free cells, moving the population into the *ghost* state (orange), and extending the half-life of plasmid decay (right). **(b – d).** Time series of average plasmid abundance (mean ± SE, n = 3) for non-mobilizable plasmids **(b)** pSC101, **(c)** colE1, and **(d)** pUC under different selection strength. Between day 2 and 3 (grey), antibiotics were applied at different concentrations to induce varying levels of ghost effect. Three representative conditions are shown here, corresponding to no antibiotic treatment (circle), 1/32 × 50 μg/mL (triangle), and 50 μg/mL (square). pSC101 and colE1 populations were treated with Kan, and pUC populations were treated with Spec. **(e – g).** Dose-dependence of plasmid half-lives τ_1/2_ (mean ± SE, delta method) on the pulsed antibiotic concentration for **(e)** pSC101, **(f)** colE1, and **(g)** pUC. Right-censored datapoints (τ_1/2_ > 17 days) are shown without error bar.

### Antibiotic pulse induces prolonged plasmid carriage in synthetic *E. coli* communities

To determine whether the ghost effect occurs in more complex microbial systems, we constructed five synthetic communities of *E. coli* Keio strains^39^ (Table S1 & S2). Each strain contains a uniquely barcoded, non-mobilizable plasmid, enabling the quantification of community structure using next generation sequencing (NGS). The first community, Comm87, consisted of 86 background strains plus one donor strain carrying conjugative plasmid R388 (trimethoprim resistant, Trim^R^). Four other communities (Comm57) shared the same background community of 56 barcoded Keio strains, in addition to a unique donor strain carrying one additional conjugative plasmid, either R6K (streptomycin resistant, Strp^R^), pCU1 (carbenicillin resistant, Carb^R^), R388, or RP4 (tetracycline resistant, Tet^R^). The overnight cultures of the background strains were equally combined, then mixed with the donor strain at a 3:1 ratio to assemble each of the five communities. The communities were passaged daily with a 100-fold dilution in LB media supplemented with 100 μg/mL Carb for Comm87 or 25 μg/mL Cm for Comm57 to maintain the barcode plasmids. We tracked the abundance of the target plasmids over time through selective plating.

Without any additional antibiotic intervention, the conjugative plasmids exponentially decayed (**Fig. 3a - c**, purple), with half-lives of 2.2 ± 0.4 days (mean ± SD, n = 3) for R388 in Comm87, 4.4 ± 0.1 days (mean ± SD, n = 3) for R6K in Comm57, and 1.0 ± 0.1 days (mean ± SD, n = 3) for pCU1 in Comm57 (**Fig. 3d - f**). R388 and RP4 were able to persist in Comm57 for the entire course of the 20-day experiment (**Fig. 3g & h**). As predicted by the model, antibiotic selection for the plasmids, applied between day 2 and 3, increased plasmid abundances to approach 100%. Driving the communities into the *ghost* state drastically extended plasmid half-lives to 27.3 ± 1.7 days (mean ± SD, n = 3) for R388 in Comm87 and 9.9 ± 0.9 days (mean ± SD, n = 3) for pCU1 in Comm57 (**Fig. 3 d-f,** p<0.01 or p<0.001 with Welch’s t-test). Plasmids R6K, R388, and RP4 all persisted in the communities at ∼100% for more than 17 days after antibiotic selection (**Fig. 3b, g, h**, orange). The four Comm57 communities, when passaged in a different dilution rate (1:500), showed similar results (**Fig. S4**).

**Figure 3.**
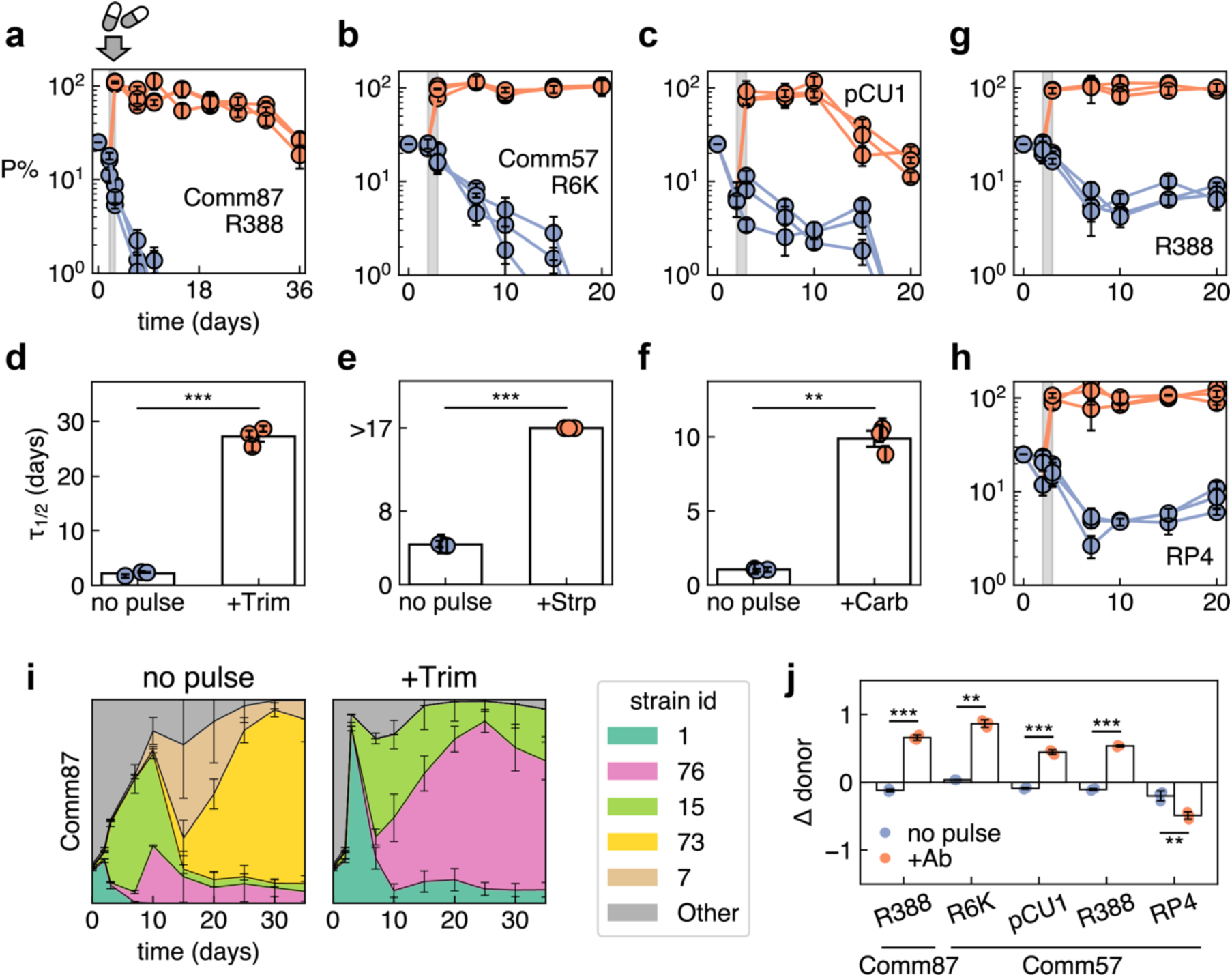
Antibiotic pulse induces prolonged plasmid carriage in synthetic *E. coli* communities. **(a – c).** Plasmid abundance (mean ± SE based on technical triplicates of selective plating) over time for **(a)** R388 in Comm87, **(b)** R6K in Comm60, **(c)** pCU1 in Comm 60. Purple curves: communities were passaged daily at 1:100 dilution ratio in LB supplemented with 100 μg/mL Carbenicillin (Comm87, **a**) or 25μg/mL Chloramphenicol (Comm57, **b & c**). Orange curves: communities were treated with an additional antibiotic pulse between day 2 and 3 (grey bar) to select for the plasmid-carrying populations (R388: 10 μg/mL Trimethoprim, R6K: 100 μg/mL Streptomycin, pCU1: 100 μg/mL Carbenicillin). **(d - f)**. Antibiotic pulses extended plasmid half-lives (mean ± SE, delta method) for **(d)** R388 in Comm87, **(e)** R6K in Comm57, **(f)** pCU1 in Comm57. Right-censored datapoints (τ_1/2_ > 17 days) are shown without error bar. For hypothesis testing, half-lives for R6K in the pulsed group (+Strp) were taken as τ_1/2_ = 17 days. **(d)** Average half-life of R388 after Trim treatment was significantly higher than in LB group: 27.3 ± 1.7 (SD, n = 3) vs 2.2 ± 0.4 (SD, n = 3) days. Welch’s t-test: t(2.21) = 24.91, p<0.001, mean difference = 25.1 [95% CI: 21.1, 29.1], Cohen’s d = 20.34. **(e)** Average half-life of R6K after Trim treatment was significantly higher than in LB group: 17.0 ± 0.0 (SD, n = 3) vs 4.4 ± 0.1 (SD, n = 3) days. Welch’s t-test: t(2.00) = 190.56, p<0.001, mean difference = 12.6 [95% CI: 12.4, 12.9], Cohen’s d = 155.59. **(f)** Average half-life of pCU1 after Trim treatment was significantly higher than in LB group: 9.9 ± 0.9 (SD, n = 3) vs 1.0 ± 0.1 (SD, n = 3) days. Welch’s t-test: t(2.01) = 16.50, p=0.004, mean difference = 8.8 [95% CI: 6.5, 11.1], Cohen’s d = 13.47. In the figures, **: p < 0.01, ***: p < 0.001. **(g & h).** Plasmid abundance (mean ± SE based on technical triplicates of selective plating) over time for **(g)** R388 and **(h)** RP4 in Comm60. Antibiotic pulsing induced alternative steady state for plasmid persistence. Purple curves: communities passaged daily at 1:100 dilution ratio in LB supplemented with 25μg/mL Chloramphenicol. Orange: communities were treated with additional antibiotic pulse between Day 2 and 3 (grey bar) to select for the plasmid-carrying populations (R388: 10 μg/mL Trimethoprim, RP4: 10 μg/mL Tetracycline). **(i).** Strain-level community dynamics (mean ± SE, n = 3) of Comm87 without (left) or with (right) 1-day of Trimethoprim treatment. The five most abundant strains (based on cumulative relative abundance across samples) are color-coded. Strain 1 corresponds to the donor strain carrying plasmid R388. **(j).** Change in the relative abundance of the donor strain (Δ donor) before (day 2) and after (day 3) the antibiotic pulse. Bars represent mean ± SE (n = 3). Welch’s t-test, **: p < 0.01, ***: p < 0.001.

We used NGS to quantify the community composition over time (**Fig. 3i & Fig. S5**). Upon antibiotic selection, a significant shift in each community structure was observed, and the community diversity dropped significantly (**Fig. S6**). Upon selection, the relative abundances of the plasmid donors were significantly amplified (**Fig. 3j**), while the donor abundances decreased in most communities without antibiotic pulse. The decline in diversity, and the increase in donor abundance, suggests that the plasmids have not yet spread to every member in the community. One exception was Comm57 harboring RP4, where the donor abundance was significantly decreased upon antibiotic selection (**Fig. 3j**). While in other communities the antibiotic pulse resulted in a crash in diversity, the RP4 community instead increased its diversity (**Fig. S6**). This could be due to the combination of two factors: the low fitness of the donor strain, and the fast conjugation capacity of RP4^24,40^. By the time positive selection took place, most strains in the community would have already received the plasmid and could outcompete the donor strain in the presence of Tet.

These findings demonstrate that the ghost effect operates not only in clonal populations but also in synthetic microbial communities. A brief antibiotic pulse can reorganize community composition by selectively eliminating plasmid-free competitors, thereby stabilizing plasmid persistence at the community level without requiring genetic adaptation.

### Ghost effect in multi-specific communities with niche partition

In nature, microbes can co-exist in natural communities through niche partition, where different members can take up different resources (**Fig. 4a**) or habitats (**Fig. 4b**) to avoid competitive exclusion. In the above experiments, we showed the ghost effect taking place in clonal populations and *E. coli* communities, where strong competitions exist between individual strains, and plasmid-free populations are sensitive to the antibiotics. In these cases, ghost effect takes place when the plasmid abundance reaches ∼100%. However, numerical simulations (**Methods**) suggest antibiotic-induced ghost effect can happen without community purity (**Fig. 4 c & d**) in the presence of niche partition. For instance, consider a mixed population of plasmid-free strain 1, plasmid-carrying strain 1, and non-host strain 2 (**Methods**). In the simple case where (1) strain 1 and 2 do not interact and (2) strain 2 has moderate resistance to the antibiotic, the non-host strain 2 can survive the antibiotic selection, limiting the plasmid abundance to < 100%. However, ghost effect can still happen within the purified strain 1 population, resulting in the prolonged carriage of the plasmid in the two-member community (**Fig. 4c & d**). In essence, as long as the plasmid-carrying population is approaching purity in a niche, ghost effect should take place. Since natural microbiomes often have niche partition, with different species exhibiting natural or acquired resistance, we expect general applicability of ghost effect following antibiotic treatments, even when the plasmid or resistance abundance does not reach 100%.

**Figure 4.**
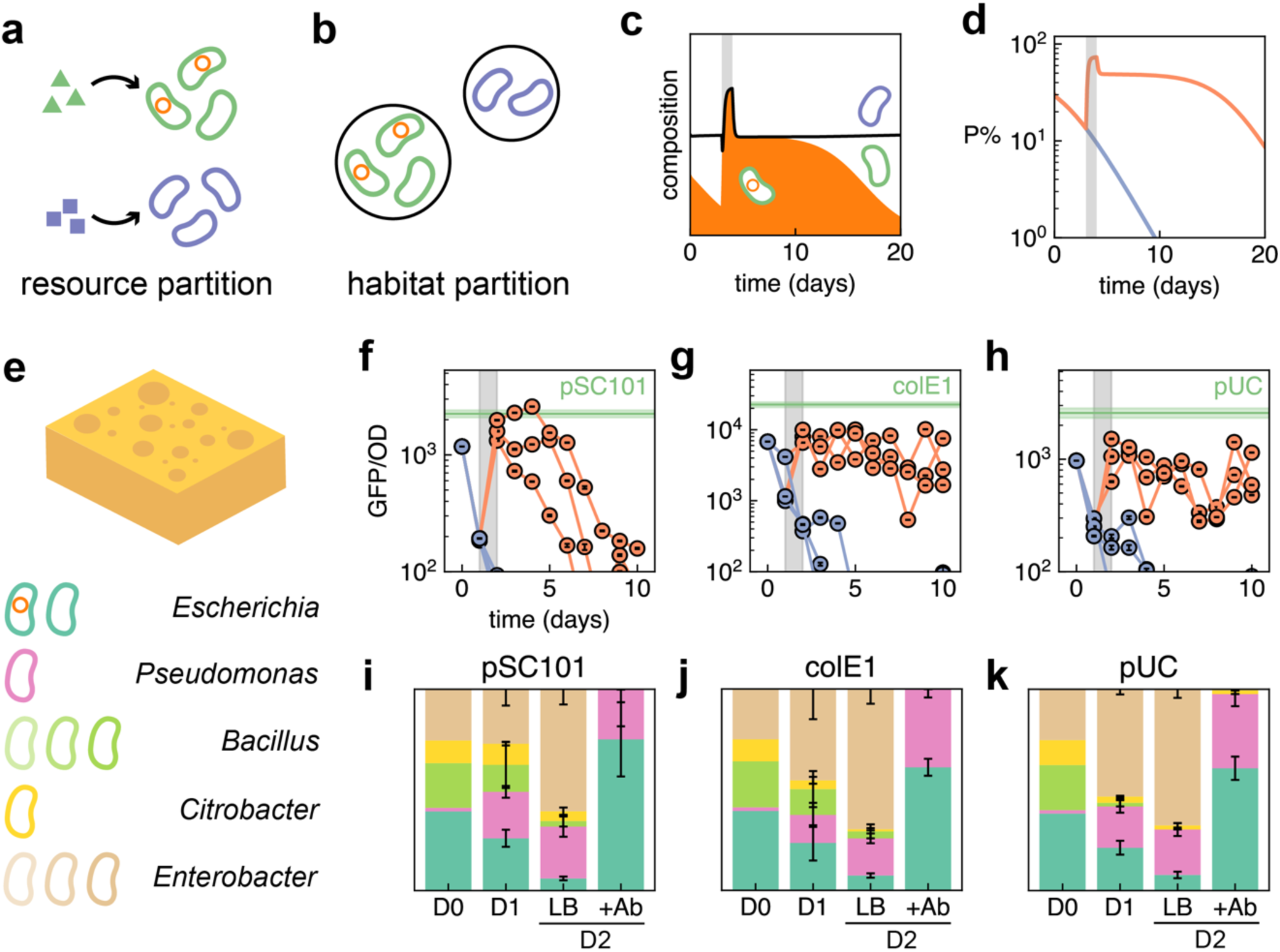
Ghost effect in niche partitioned communities. **(a & b).** Microbes co-exist through partitioning of resource or habitat. **(c).** Change in the community composition over time for a mixed population of two strains (green and purple) with complete niche partitioning (no interaction). Orange shading indicates the plasmid-carrying fraction of strain 1 (green). The community experienced selection for the plasmid-carrying cells between day 2 and 3 (gray bar). **(d).** Plasmid abundance over time with (orange) and without (purple) 1-day antibiotic selection (gray bar) for the plasmid. The ghost effect occurs without plasmid abundance reaching P% = 100%. **(e).** Microbial communities containing plasmid-free and plasmid-carrying *E. coli* MG1655 and eight sink isolates were cultured in LB media with sponge-imposed structure. Genus-level taxonomies of the eight sink isolates are shown. The isolates include one *Pseudomonas* strain (*P. aeruginosa*), three *Bacillus* strains, one *Citrobacter* strain, and three *Enterobacter* strains (including one strain identified as *E. cloacae*). Strains are color-coded by genus correspondingly as in (**i – k**). Three plasmids (pSC101, colE1, and pUC) were separately cultured in three independent communities. **(f – h).** Time series of GFP/OD for GFP-encoding plasmids pSC101, colE1, and pUC. Purple: communities were passaged daily at 1:2×10^7^ dilution ratio in sponges immersed in LB. Orange: communities were treated with additional antibiotic pulse between day 1 and 2 (grey bar) to select for the plasmid-carrying populations (pSC101 and colE1: 50 μg/mL Kanamycin, pUC: 50 μg/mL Spectinomycin). Green bar: mean ± SE GFP/OD for *E. coli* MG1655 carrying the corresponding plasmid (n = 3). Error bars indicate propagated measurement uncertainties (per-well SD). **(i – k).** 16S-resolved community dynamics between day 0 (D0) and day 2 (D2) (n = 1 for day 0, mean ± SE, n = 3 for other samples). After the antibiotic pulse, the communities were dominated by *E. coli* and *P. aeruginosa*. Due to limited taxonomic resolution of 16S sequencing, *E. coli* was assigned as *Escherichia-Shigella*, here we denote it as *Escherichia*. Taxa with maximum relative abundance < 1% across all samples are not shown.

To test this idea, we constructed three communities consisting of 8 isolates from Duke Hospital sink p-traps^41^ (**Methods**), plasmid-free and plasmid-carrying *E. coli* MG1655 with one of the three non-mobilizable plasmids (pSC101, colE1 and pUC) expressing sfGFP as a surrogate for their abundance. The sink isolates consist of one *Pseudomonas aeruginosa* strain, three *Bacillus* trains, one *Citrobacter* strain, and three *Enterobacter* strains (**Fig. 4e**). To facilitate niche partitioning, we used cellulose sponges immersed in LB media to create porous, physically partitioned habitats (**Fig. 4e**), which were previously found to promote the diversity of microbial communities^42^.

In the absence of antibiotic selection, the GFP/OD level (**Methods**) dropped rapidly to below 10^2^. After transient Kan or Spec selection between day 1 and 2, GFP levels were increased. pSC101 showed slowed decay in GFP/OD level, while colE1 and pUC persisted until day 10 (**Fig. 4 f – h**). This is consistent with our observations of the three plasmids in clonal populations (**Fig. 1, Fig. S1**). Importantly, for plasmids colE1 and pUC, the GFP/OD levels were consistently below the baseline in pure plasmid-carrying MG1655 populations.

Through 16S sequencing (**Methods**), we quantified community dynamics in the presence and absence of the transient antibiotic treatment (**Fig. 4 i – k**). On day 2, in the absence of antibiotics, *P. aeruginosa* and *Enterobacter* became dominant in the communities, with the other three genera decreasing in their fractions or even become undetectable in some of the replicates. Antibiotic treatment increased the abundance of the two resistant strains, *E. coli* and *P. aeruginosa*, and eliminated the rest of the community. The survival of *P. aeruginosa* confirms that the communities do not purely contain plasmid-carrying cells, but a ghost effect within the *E. coli* population still takes place. Consistent with the GFP/OD reading of pSC101, *P. aeruginosa* was detectable in one of the three replicate communities harboring pSC101, potentially due to the stochastic effect of high dilution rate (2×10^7^) during daily passaging.

In LB cultures without cellulose sponges, the GFP/OD levels of the communities after antibiotic treatment became comparable to the pure plasmid-carrying baselines (Fig. S7) and exhibited slowed-down decay in comparison to the antibiotic-free control group. Since the plasmids are non-mobilizable, the post-treatment communities likely consisted primarily of *E. coli*. In contrast, the survival of *P. aeruginosa* should be attributed to the presence of sponges, which possibly introduces spatial structures to mediate the competition^42^ and promote biofilm formation^43^.

## Discussion

Resistance limits the clinical efficacy of antibiotic treatments and poses a major threat to human health. Many antibiotic resistance genes are maintained and propagated by mobile genetic elements including plasmids^1–3^. Understanding and predicting the persistence of plasmids is critical for designing intervention strategies to contain and reverse the spread of AMR. In this work, we systematically quantified the decay dynamics of multiple plasmids across different community contexts, and established their dependence on community structure. We showed that the plasmid half-life drastically increases in nearly pure plasmid-carrying populations, either from assembly or induced by transient antibiotic pulse. Theoretically, this ghost effect arises from the decrease in the strength of growth competition when plasmid-free cells are scarce in the population.

Our results reveal a common mechanism underlying slow reversal of plasmid-mediated resistance after antibiotic treatments, a phenomenon that spans across microbial ecosystems ranging from the human oral, urinary, and gut microbiomes, to those of livestock. Macrolide antibiotics have been reported to increase the abundance of resistant streptococci (primarily carrying *mef* gene or macrolide-lincosamide-streptogramin-B resistant cassette *erm*(B)) in oral microbiomes to more than 80%, which required over 6 months to revert to baseline^9^. The authors attributed this prolonged resistance carriage to the long half-life of azithromycin, and the low burden of the *erm*(B) gene. Similarly, veterinary β-lactams strongly amplified the abundance of resistant bacteria in the pig intestinal microbiome, persisting for more than 3 weeks beyond the drug withdrawal. The extended-spectrum β-lactamase CTX-M-1 was carried by both an exogenous conjugative plasmid and indigenous gut microbiota. Consistent with our results (**Fig. 3, Fig. S5**), the authors observed the significant amplification of resistance abundance and donor abundance upon antibiotic selection, followed by rapid decline of the donor strain abundance after antibiotic withdrawal. Importantly, both the *erm*(B) cassette ^13–17^ and the CTX-M-1 gene^18,19^ are commonly found on conjugative plasmids and other mobilizable gene elements. Yelin et al. reported that resistance in urinary tract infections shows autocorrelation lasting more than six months, with strong and persistent correlation between patients’ antibiotic purchase history and subsequent resistance^12^. Anthony et al. observed increases in the abundance of β-lactam, macrolide, and tetracycline resistance following treatment with cephalosporin (β-lactam), azithromycin (macrolide), or both^10^. During the development of the infant gut microbiome, resistance genes on mobile genetic elements were found to persist longer than those encoded on chromosomes^11^. The diversity of antibiotics, resistance genes, bacterial host, and ecosystems all point to a general population-dynamical mechanism. As the ghost effect depends only on the slow time scale of segregation loss, it offers a potential explanation for these observations.

While our work focuses on the population dynamics of the plasmids, the ghost effect also has strong evolutionary consequences. Prolonged plasmid carriage enables co-evolution of the plasmid and host, creating opportunities for compensatory mutations. Long-term co-evolution could ameliorate the burden of the plasmid^20^, and may even turn the deleterious cost of plasmid carriage to a beneficial one^22^. Thus, despite predictions that certain plasmids should not persist based on ecological models^24,27^, their long-term persistence may occur before eventual extinction. In our experiments, we observed potential evolutionary rescues for colE1 and pUC starting at high initial plasmid abundances (**Fig. S1**), which was not observed in populations outside of the ghost effect region. Therefore, plasmid persistence can have bistable outcomes when evolution forces are taken into consideration.

The antibiotic-induced persistence also provides an environment-driven explanation for the survival of burdensome plasmids, addressing the so-called plasmid paradox^7^. Previous studies have attributed plasmid survival to biotic factors, such as fast conjugation^24,25,44^, variable fitness effects across different hosts^23^, spatial partitioning^45^, and compensatory mutations that ameliorates the plasmid burden^20,22^. These works typically assume constant environments, without positive selections on the plasmids. However, plasmids in natural and clinical settings experience fluctuating selection forces^46–50^. In such cases, the burdensome plasmids can persist when the frequency of positive selection is on the same order of magnitude as the plasmid half-life within the ghost effect region. This mechanism further raises questions about the efficacy of simply reducing antibiotic use to contain AMR^6^ – similar persistence can be achieved with far less amount of antibiotics with sporadic selection. Conversely, pulsed antibiotic treatments has been proposed as a strategy to constrain the development of resistance^51–55^. Such strategies should be designed to avoid the ghost effect region to prevent slow population responses that disproportionally favor resistant bacteria.

The clinical, ecological, and evolutionary implications of ghost effect underscore the need for strategies that counteract against unintended antibiotic exposure. Previous studies have used chemicals to reduce plasmid abundance and reverse its persistence^24,56^. As both plasmid abundance and half-life are governed by conjugation, growth, and segregation events^27^, we hypothesize that these chemicals could also accelerate plasmid decay. We found that curing chemicals and their combinations were able to reduce the half-life of pSC101 almost by half (**Fig. S8**). A single day of treatment using chemical combinations achieved comparable effects as full course, suggesting these chemicals may accelerate the plasmid decay by increasing segregation error.

Overall, our work established the timescale decoupling between segregation loss and growth competition for burdensome plasmids and demonstrated the disproportionate impact of transient antibiotic pulses on the plasmid persistence. Our findings provide a generalizable mechanism for the prolonged ecological memory of plasmid carriage following antibiotic exposure, and have implications for clinical management of antibiotic use and the ecology and evolution of plasmids.

## Acknowledgments

We thank Emma Chory, Teng Wang, Rohan Maddemsetti, Emrah Şimşek, Grayson Hamrick, Yifan Dai, and Aaron Yip for helpful discussions. This work was supported by National Institute of Health (L.Y.: R01AI125604, R01GM098642, R01EB031869) and the National Science Foundation (L. Y.: Cooperative Agreement No. EEC-2133504)

## Author contributions

Z.Z., A.W., and L.Y. conceived the research. Z.Z., A.W., Z.Y., X.C., and K.L. performed the experiments. Z.Z. performed the modelling and statistical analysis. Z.Z., A.W., and L.Y. wrote the manuscript. All authors read and contributed to the manuscript.

## Declaration of generative AI and AI-assisted technologies in the writing process

During the preparation of this work the authors used ChatGPT to improve the language of the manuscript. After using this tool, the authors reviewed and edited the content as needed and take full responsibility for the content of the published article.

## Resource availability

### Materials availability

Genetic constructs generated for this study are available upon request.

### Lead contact

Further information and requests for resources should be directed to and will be fulfilled by the lead contact, Lingchong You.

### Data and code availability

All original code and data have been deposited at https://github.com/youlab/Plasmid-Ghost-Effect.

## References

1 Bethke, J. H. et al. Environmental and genetic determinants of plasmid mobility in pathogenic *Escherichia coli*. Science Advances 6, eaax3173 (2020). 10.1126/sciadv.aax3173

2 Lipworth, S. et al. The plasmidome associated with Gram-negative bloodstream infections: A large-scale observational study using complete plasmid assemblies. Nature Communications 15 (2024). 10.1038/s41467-024-45761-7

3 Castañeda-Barba, S., Top, E. M. & Stalder, T. Plasmids, a molecular cornerstone of antimicrobial resistance in the One Health era. Nature Reviews Microbiology 22, 18–32 (2024). 10.1038/s41579-023-00926-x

4 Hawkey, P. M. & Jones, A. M. The changing epidemiology of resistance. J Antimicrob Chemother 64 **Suppl 1**, i3–10 (2009). 10.1093/jac/dkp256

5 Mathers, A. J., Peirano, G. & Pitout, J. D. The role of epidemic resistance plasmids and international high-risk clones in the spread of multidrug-resistant Enterobacteriaceae. Clin Microbiol Rev 28, 565–591 (2015). 10.1128/cmr.00116-14

6 Andersson, D. I. & Hughes, D. Antibiotic resistance and its cost: is it possible to reverse resistance? Nature Reviews Microbiology 8, 260–271 (2010). 10.1038/nrmicro2319

7 Brockhurst, M. A. & Harrison, E. Ecological and evolutionary solutions to the plasmid paradox. Trends in Microbiology 30, 534–543 (2022). 10.1016/j.tim.2021.11.001

8 Cavaco, L. M., Abatih, E., Aarestrup, F. M. & Guardabassi, L. Selection and persistence of CTX-M-producing Escherichia coli in the intestinal flora of pigs treated with amoxicillin, ceftiofur, or cefquinome. Antimicrob Agents Chemother 52, 3612–3616 (2008). 10.1128/aac.00354-08

9 Malhotra-Kumar, S., Lammens, C., Coenen, S., Van Herck, K. & Goossens, H. Effect of azithromycin and clarithromycin therapy on pharyngeal carriage of macrolide-resistant streptococci in healthy volunteers: a randomised, double-blind, placebo-controlled study. The Lancet 369, 482–490 (2007). 10.1016/S0140-6736(07)60235-9

10 Anthony, W. E. et al. Acute and persistent effects of commonly used antibiotics on the gut microbiome and resistome in healthy adults. Cell Reports 39, 110649 (2022). 10.1016/j.celrep.2022.110649

11 Yassour, M. et al. Natural history of the infant gut microbiome and impact of antibiotic treatment on bacterial strain diversity and stability. Sci Transl Med 8, 343ra381 (2016). 10.1126/scitranslmed.aad0917

12 Yelin, I. et al. Personal clinical history predicts antibiotic resistance of urinary tract infections. Nature Medicine 25, 1143–1152 (2019). 10.1038/s41591-019-0503-6

13 Berryman, D. I. & Rood, J. I. The closely related ermB-ermAM genes from Clostridium perfringens, Enterococcus faecalis (pAM beta 1), and Streptococcus agalactiae (pIP501) are flanked by variants of a directly repeated sequence. Antimicrob Agents Chemother 39, 1830–1834 (1995). 10.1128/aac.39.8.1830

14 Clewell, D. B., Flannagan, S. E. & Jaworski, D. D. Unconstrained bacterial promiscuity: the Tn916-Tn1545 family of conjugative transposons. Trends Microbiol 3, 229–236 (1995). 10.1016/s0966-842x(00)88930-1

15 Horodniceanu, T., Bouanchaud, D. H., Bieth, G. & Chabbert, Y. A. R plasmids in Streptococcus agalactiae (group B). Antimicrob Agents Chemother 10, 795–801 (1976). 10.1128/aac.10.5.795

16 Leblanc, D. J. & Lee, L. N. Physical and genetic analyses of streptococcal plasmid pAM beta 1 and cloning of its replication region. Journal of Bacteriology 157, 445–453 (1984). 10.1128/jb.157.2.445-453.1984

17 Shaw, J. H. & Clewell, D. B. Complete nucleotide sequence of macrolide-lincosamide-streptogramin B-resistance transposon Tn917 in Streptococcus faecalis. J Bacteriol 164, 782–796 (1985). 10.1128/jb.164.2.782-796.1985

18 Cormier, A. C. et al. Diversity of blaCTX-M-1-carrying plasmids recovered from Escherichia coli isolated from Canadian domestic animals. PLOS ONE 17, e0264439 (2022). 10.1371/journal.pone.0264439

19 Dolejska, M., Villa, L., Hasman, H., Hansen, L. & Carattoli, A. Characterization of IncN plasmids carrying bla CTX-M-1 and qnr genes in Escherichia coli and Salmonella from animals, the environment and humans. J Antimicrob Chemother 68, 333–339 (2013). 10.1093/jac/dks387

20 San Millan, A., et al. Positive selection and compensatory adaptation interact to stabilize non-transmissible plasmids. Nature Communications 5, 5208 (2014). 10.1038/ncomms6208

21 Enne, V. I. et al. Assessment of the fitness impacts on Escherichia coli of acquisition of antibiotic resistance genes encoded by different types of genetic element. Journal of Antimicrobial Chemotherapy 56, 544–551 (2005). 10.1093/jac/dki255

22 Loftie-Eaton, W. et al. Compensatory mutations improve general permissiveness to antibiotic resistance plasmids. Nature Ecology & Evolution 1, 1354–1363 (2017). 10.1038/s41559-017-0243-2

23 Alonso-del Valle, A., et al. Variability of plasmid fitness effects contributes to plasmid persistence in bacterial communities. Nature Communications 12, 2653 (2021). 10.1038/s41467-021-22849-y

24 Lopatkin, A. J. et al. Persistence and reversal of plasmid-mediated antibiotic resistance. Nature Communications 8 (2017). 10.1038/s41467-017-01532-1

25 Hall, J. P. J., Wood, A. J., Harrison, E. & Brockhurst, M. A. Source–sink plasmid transfer dynamics maintain gene mobility in soil bacterial communities. Proceedings of the National Academy of Sciences 113, 8260–8265 (2016). 10.1073/pnas.1600974113

26 Alonso-Del Valle, A., et al. Antimicrobial resistance level and conjugation permissiveness shape plasmid distribution in clinical enterobacteria. Proceedings of the National Academy of Sciences 120 (2023). 10.1073/pnas.2314135120

27 Wang, T. & You, L. The persistence potential of transferable plasmids. Nature Communications 11 (2020). 10.1038/s41467-020-19368-7

28 Wang, X. et al. Inter-plasmid transfer of antibiotic resistance genes accelerates antibiotic resistance in bacterial pathogens. Isme j 18 (2024). 10.1093/ismejo/wrad032

29 Murray, L. M. et al. Co-selection for antibiotic resistance by environmental contaminants. npj Antimicrobials and Resistance 2, 9 (2024). 10.1038/s44259-024-00026-7

30 Cooper, N. S., Brown, M. E. & Caulcott, C. A. A Mathematical Method for Analysing Plasmid Stability in Micro-organisms. Microbiology 133, 1871–1880 (1987). 10.1099/00221287-133-7-1871

31 Summers, D. K. The kinetics of plasmid loss. Trends in Biotechnology 9, 273–278 (1991). 10.1016/0167-7799(91)90089-Z

32 Fedorec, A. J. H. et al. Two New Plasmid Post-segregational Killing Mechanisms for the Implementation of Synthetic Gene Networks in Escherichia coli. iScience 14, 323–334 (2019). 10.1016/j.isci.2019.03.019

33 Danino, T. et al. Programmable probiotics for detection of cancer in urine. Science Translational Medicine 7, 289ra284–289ra284 (2015). 10.1126/scitranslmed.aaa3519

34 Nordström, K. & Austin, S. J. MECHANISMS THAT CONTRIBUTE TO THE STABLE SEGREGATION OF PLASMIDS. Annual Review of Genetics 23, 37–69 (1989). 10.1146/annurev.ge.23.120189.000345

35 Baxter, J. C. & Funnell, B. E. Plasmid Partition Mechanisms. Microbiol Spectr 2 (2014). 10.1128/microbiolspec.PLAS-0023-2014

36 Cooper, T. F. & Heinemann, J. A. Postsegregational killing does not increase plasmid stability but acts to mediate the exclusion of competing plasmids. Proceedings of the National Academy of Sciences 97, 12643–12648 (2000). 10.1073/pnas.220077897

37 Strogatz, S. a. Nonlinear dynamics and chaos : with applications to physics, biology, chemistry, and engineering. (Second edition. Boulder, CO : Westview Press, a member of the Perseus Books Group, [2015], 2015).

38 Shao, B. et al. Single-cell measurement of plasmid copy number and promoter activity. Nature Communications 12, 1475 (2021). 10.1038/s41467-021-21734-y

39 Baba, T. et al. Construction of Escherichia coli K-12 in-frame, single-gene knockout mutants: the Keio collection. Mol Syst Biol 2, 2006.0008 (2006). 10.1038/msb4100050

40 Wang, T. et al. Horizontal gene transfer enables programmable gene stability in synthetic microbiota. Nature Chemical Biology (2022). 10.1038/s41589-022-01114-3

41 Şimşek, E., et al. Keystone engineering enables collective range expansion in microbial communities. bioRxiv, 2025.2001.2011.632568 (2025). 10.1101/2025.01.11.632568

42 Wu, F. et al. Modulation of microbial community dynamics by spatial partitioning. Nature Chemical Biology 18, 394–402 (2022). 10.1038/s41589-021-00961-w

43 Agustín, M. D., Stengel, P., Kellermeier, M., Tücking, K.-S. & Müller, M. Monitoring Growth and Removal of Pseudomonas Biofilms on Cellulose-Based Fabrics. Microorganisms 11 (2023).

44 Stevenson, C., Hall, J. P. J., Harrison, E., Wood, A. J. & Brockhurst, M. A. Gene mobility promotes the spread of resistance in bacterial populations. The ISME Journal 11, 1930–1932 (2017). 10.1038/ismej.2017.42

45 Weiss, A., Wang, T. & You, L. Promotion of plasmid maintenance by heterogeneous partitioning of microbial communities. Cell Systems 14, 895–905.e895 (2023). 10.1016/j.cels.2023.09.002

46 Stevenson, C., Hall, J. P. J., Brockhurst, M. A. & Harrison, E. Plasmid stability is enhanced by higher-frequency pulses of positive selection. Proceedings of the Royal Society B: Biological Sciences 285, 20172497 (2018). 10.1098/rspb.2017.2497

47 Cai, M. et al. Occurrence and temporal variation of antibiotics and antibiotic resistance genes in hospital inpatient department wastewater: Impacts of daily schedule of inpatients and wastewater treatment process. Chemosphere 292, 133405 (2022). 10.1016/j.chemosphere.2021.133405

48 Cao, S., Zhang, P., Halsall, C., Hou, Z. & Ge, L. Occurrence and seasonal variations of antibiotic micro-pollutants in the Wei River, China. Environmental Research 252, 118863 (2024). 10.1016/j.envres.2024.118863

49 Schuster, D. et al. Antibiotic concentrations in raw hospital wastewater surpass minimal selective and minimum inhibitory concentrations of resistant Acinetobacter baylyi strains. Environ Microbiol 24, 5721–5733 (2022). 10.1111/1462-2920.16206

50 Coutu, S., Rossi, L., Barry, D. A., Rudaz, S. & Vernaz, N. Temporal Variability of Antibiotics Fluxes in Wastewater and Contribution from Hospitals. PLOS ONE 8, e53592 (2013). 10.1371/journal.pone.0053592

51 Baker, C. M., Ferrari, M. J. & Shea, K. Beyond dose: Pulsed antibiotic treatment schedules can maintain individual benefit while reducing resistance. Scientific Reports 8, 5866 (2018). 10.1038/s41598-018-24006-w

52 Letten, A. D., Hall, A. R. & Levine, J. M. Using ecological coexistence theory to understand antibiotic resistance and microbial competition. Nature Ecology & Evolution 5, 431–441 (2021). 10.1038/s41559-020-01385-w

53 Bauer, M., Graf, I. R., Ngampruetikorn, V., Stephens, G. J. & Frey, E. Exploiting ecology in drug pulse sequences in favour of population reduction. PLOS Computational Biology 13, e1005747 (2017). 10.1371/journal.pcbi.1005747

54 Lin, W.-H. & Kussell, E. Complex Interplay of Physiology and Selection in the Emergence of Antibiotic Resistance. Current Biology 26, 1486–1493 (2016). 10.1016/j.cub.2016.04.015

55 Nev, O. A., Jepson, A., Beardmore, R. E. & Gudelj, I. Predicting community dynamics of antibiotic-sensitive and -resistant species in fluctuating environments. Journal of The Royal Society Interface 17, 20190776 (2020). 10.1098/rsif.2019.0776

56 Buckner, M. M. C., Ciusa, M. L. & Piddock, L. J. V. Strategies to combat antimicrobial resistance: anti-plasmid and plasmid curing. FEMS Microbiology Reviews 42, 781–804 (2018). 10.1093/femsre/fuy031

