## Supplemental Material for "Dynamical memory underlies prolonged plasmid persistence after transient antibiotic treatment"

This PDF includes:

- Methods
- Figure S1 – S8
- Table S1 & S2
- Supplementary note

### Methods

#### Mathematical modelling of plasmid dynamics

To investigate the plasmid dynamics in the clonal population, a previous model describing the conjugative plasmid dynamics was used.[1]

$$\frac{df}{dt} = \alpha\mu\left(1 - \frac{f+p}{c}\right)f - Df + \kappa p - \eta f p$$

$$\frac{dp}{dt} = \mu\left(1 - \frac{f+p}{c}\right)p - Dp - \kappa p + \eta f p$$

Here $f$ denotes the cell density of plasmid-free cells, $p$ denotes the cell density of plasmid-carrying cells. $\mu$ is the growth rate of plasmid-carrying cells, $\alpha$ reflects the fitness effect of the plasmid: costly when $\alpha$ > 1 or beneficial when $\alpha$ <1, $c$ is the carrying capacity, $D$ is the dilution rate, $\kappa$ is the segregation loss rate, and $\eta$ is the conjugation efficiency of the plasmid from plasmid-carrying cells to plasmid-free cells.

Cooper et al.[2] obtained an analytical solution of non-mobilizable plasmid dynamics by assuming constant difference in the growth rates between plasmid-free and plasmid-carrying cells. Similar to this derivation, we extended the analytical solution to conjugative plasmid (**Supplementary note**). The temporal dynamics of plasmid abundance (fraction of plasmid-carrying cells) $P\% = p/(f+p)$ can be expressed as

$$P\%(t) = \frac{aP_0\%}{(a - bP_0\%)\exp(at) + bP_0\%},$$

where

$$a = \delta + \kappa - \eta, b = \delta - \eta$$

$$\delta = (\alpha - 1)\mu\left(1 - \frac{f + p}{c}\right).$$

As the total cell density $f + p$ is not a constant, we can show

$$\left(1 - \frac{1}{\alpha}\right)D \le \delta \le (\alpha - 1)D.$$

The plasmid half-life can then be derived as

$$\tau_{1/2} = \frac{1}{a}\log\left(\frac{2a - bP_0}{a - bP_0}\right) \quad (1)$$

To simulate the burdensome plasmid dynamics in clonal populations, we used the following parameters: $\mu = 0.6\ hr^{-1}$, $\eta = 0$, $\kappa = 5 \times 10^{-5}\ hr^{-1}$, $D = 0.2\ hr^{-1}$, $\alpha = 1.1$, $c = 1$. Simulations were performed with initial population sizes of $p_0 = 1, 0.99, 0.9, 0.5, 0.2$, and $f_0 = 1 - p_0$, using Python scipy.integrate.odeint package with default parameters and time step of $0.1\ hr$.

As discussed in **Supplementary Notes**, $\delta$ is a variable parameter. To fit **Eq. (1)** to the simulation data in **Fig. 1c**, we used Python scipy.optimize.curve_fit for nonlinear least square fit by parameterizing $a$ and $b$ in **Eq. (1)**, with appropriate initial conditions.

To simulate the effect of antibiotics on the burdensome plasmid dynamics in communities with niche partition, we extended the model above to include an additional population. Here we denote the two populations as

$$\frac{df_1}{dt} = \alpha\mu\left(1 - \frac{s_1 + \gamma_{12}s_2}{c_1}\right)f_1 - Df_1 + \kappa p_1 - \eta f_1(p_1 + p_2) - A_1 f_1$$

$$\frac{dp_1}{dt} = \mu\left(1 - \frac{s_1 + \gamma_{12}s_2}{c_1}\right)p_1 - Dp_1 - \kappa p_1 + \eta f_1(p_1 + p_2)$$

$$\frac{df_2}{dt} = \alpha\mu\left(1 - \frac{s_2 + \gamma_{21}s_1}{c_2}\right)f_2 - Df_2 + \kappa p_2 - \eta f_2(p_1 + p_2) - A_2 f_2$$

$$\frac{dp_2}{dt} = \mu\left(1 - \frac{s_2 + \gamma_{21}s_1}{c_2}\right)p_2 - Dp_2 - \kappa p_2 + \eta f_2(p_1 + p_2)$$

Here $s_1 = f_1 + p_1$, and $s_2 = f_2 + p_2$. The key additional assumptions of this model are: 1) populations 1 and 2 engage in interactions with coefficient $\gamma_{12} < 1$ and $\gamma_{21} < 1$, and 2) the plasmid-free populations are killed by antibiotics at rate $A_1 f_1$ and $A_2 f_2$. For simplicity, we take the two populations to have the same growth rate $\mu = 0.6\ hr^{-1}$, same plasmid burden $\alpha = 1.1$, segregation loss rate $\kappa = 5 \times 10^{-5}\ hr^{-1}$, and no conjugation $\eta = 0$. We assume complete niche partition $\gamma_{12} = \gamma_{21} = 0$, and $c_1 = c_2 = 1$. Importantly, we assumed population 2 are more killed slower by the antibiotic, with $A_1 = 1hr^{-1}$ and $A_2 = 0.3hr^{-1}$. Simulation was performed with initial condition of $f_1 = 0.2$, $f_2 = 0.5$, $p_1 = 0.3$ and $p_2 = 0$. Under the conditions that population 2 is an incompatible host, and is killed slower than population 1, ghost effect can take place without the plasmid abundance reaching ~ 100%.

**Quantifying non-mobilizable plasmid dynamics in clonal populations with different initial plasmid abundances**

We started the experiment by inoculating from glycerol stock in 500 µL LB Broth (Apex LB Broth Mix, Genesee Scientific, 11-120) for 24 hours in deep well plates (Genesee, 27-413S) at 37 °C with shaking at 700 rpm and amplitude of 2mm, with corresponding selections for the plasmids: 50 µg/ml Kanamycin (Thermo Fisher, 11815032) for pSC101 and colE1, 50 µg/ml Spectinomycin (Sigma Aldrich, S4014) for pUC. 1 µL of each culture was then inoculated into 500 µL LB in deep well plates with corresponding selections. After growing for another 24 hours at 37 °C, the subsequent cultures of plasmid-carrying cells and plasmid-free cells were diluted in fresh LB to yield optical density at 600 (9) nm (OD) ≃ 0.5, mixed in different volume ratios ($P_0$% = 100%, 99%, 90%, 50%, 20%, and 0%), with OD and GFP (excitation: 488 (9) nm, emission: 520 (20) nm) measured (Tecan 200 Pro Plate Reader). The GFP/OD ratio after subtracting the baseline values of LB were linearly regressed to the plasmid abundance $P_0$%, yielding estimates of the slope (±SE) and intercept (±SE) later used to quantify the plasmid abundance in the culture based on OD and GFP measurements.

The mixed populations with different initial plasmid abundances were then cultured in LB in deep well plates, shaken with 700 rpm, 2mm amplitude, at 37 °C, and diluted 1:500 every 24 hours, for 12 days. For every 24 hours, the cultures were diluted 4-fold (50 µL of culture into 150 μL LB) to measure OD (0.4~0.6) and corresponding GFP intensity. After subtracting the baseline LB readings, the GFP/OD ratios were used to calculate the plasmid abundance. On a single day, variances in OD and GFP measurements in blank controls (pure LB) were calculated to represent the measurement uncertainty in the GFP/OD readout. This measurement uncertainty, together with the uncertainties in the estimated slope and intercept in the calibration curve, were used to calculate the per-well measurement uncertainty (SD) in plasmid-abundance through delta method.

**Quantifying non-mobilizable plasmid dynamics in clonal populations with different antibiotic pulse**

The calibration of GFP/OD reading with respect to plasmid abundance, and the initialization of the experiments is similar to described above. We started the experiment by inoculating from glycerol stock in 500 μL LB Broth for 24 hours in deep well plates at 37 °C with shaking at 700 rpm and amplitude of 2mm, with 50 μg/ml Kan for pSC101 and colE1, 50 μg/ml Spec for pUC. 1 μL of each culture was then inoculated into 500 μL LB in deep well plates with corresponding selections. After growing for another 24 hours at 37 °C, the subsequent cultures of plasmid-carrying cells and plasmid-free cells were diluted in fresh LB to OD ≃ 0.5, mixed in different volume ratios ($P_0$% = 100%, 80%, 50%, 20%, and 0%), with OD and GFP measured. The GFP/OD ratio after subtracting the baseline values of LB were linearly regressed (σ-weighted) to the plasmid abundance $P_0$%, yielding estimates of the slope *m* (±SE) and intercept *b* (±SE) later used to quantify the plasmid abundance in the culture based on OD and GFP measurements.

The equally mixed populations ($P_0$% = 50%) were then cultured in LB in deep well plates, shaken with 700 rpm, 2mm amplitude, at 37 °C, and diluted 1:500 every 24 hours. Between day 2 and 3, the populations were cultured in LB or LB with serially diluted antibiotics (with dilution rate of $2^6$, $2^5$, …, $2^0$ of 50 µg/ml Kan or 50 µg/ml Spec). After 24 hours of antibiotic pulse, all communities were transferred to LB media and cultured for another 17 days with 1:500 dilution rate. For every 24 hours, the cultures were diluted 4-fold (50 µL of culture into 150 µL LB) to measure OD (0.4~0.6) and corresponding GFP intensity. After subtracting the baseline LB readings, the GFP/OD ratios were used to calculate the plasmid abundance.

Kan (aminoglycoside) and Spec (aminocyclitol) both target the 30S ribosomal subunit and inhibit translation, thus reducing the expression of GFP. Under antibiotic selection, plasmid copy number increases, increasing plasmid encoded GFP level. We calibrated the GFP/OD to plasmid abundance using mixture standards prepared with Kan and Spec due to the potentially unknown plasmid abundance drift related to segregation error.

Calibration based on mixtures prepared without antibiotics produces the same qualitative trends, indicating our conclusions do not depend on the calibration regime.

**Quantifying conjugative plasmid dynamics in clonal populations using selective plating**

*E. coli* DA28102 cells (chloramphenicol resistant, $Cm^R$) carrying either the pCU1 (carbenicillin resistant, $Carb^R$) or R6K (streptomycin resistant, $Strp^R$) plasmid and plasmid-free DA28102 were cultured overnight in 500 µL LB supplemented with 25 µg/ml Cm (Sigma Aldrich, C0378) and appropriate antibiotics for the plasmids: 100 µg/ml Carb (Thermo Fisher, 10177012) for pCU1 and 100 µg/ml Strp (Sigma Aldrich, S6501) for R6K. The overnight cultures were diluted to OD ≃ 0.5, and plasmids-carrying cells were mixed with plasmid-free cells at different volume ratios, yielding $P_0$% of 100%, 99%, 90%, 50% or 20%. Communities were cultured at 37 °C in LB with 25 µg/ml Cm to prevent contamination, shaken at 700 rpm with amplitude of 2mm amplitude, and diluted 1:100 (5 µL into 500 µL) every 24 hours for 25 days.

The plasmid abundance was quantified using 100 µg/ml Carb or 100 µg/ml Strp selective plating (LB agar, Genesee, 11-122) for pCU1 or R6K, respectively, every 3 to 5 days using drip plating. Briefly, after making an OD measurement to estimate cell culture density, each culture was serially diluted by $10^4$ to $4×10^6$ fold, such that a 10 μL sample would contain 30-150 CFUs on LB plates. A 10 μL droplet from the diluted culture was then placed on LB or LB + selection plate and the plate was place at a 90° angle to allow droplet to run down the plate. Plates were then allowed to dry flat on the lab bench before being placed at 37 °C for colony formation. Colonies were counted and the percent plasmid carrying cells can be quantified after 24 hours. When plasmid abundance is low, multiple low dilutions of the sample were additionally plated on LB + selection plates to quantify their abundances.

**Extraction of plasmid half-lives**

The measured time-series of plasmid abundance (mean ± SD for GFP/OD measurement or SE for selective plating) from each biological replicate were log-transformed, and linearly interpolated to determine the half-life of plasmid abundance decay with respect to the initial mixture ratio. The SEs of half-life were estimated using delta method. For samples where the plasmid abundances never dropped by half, the half-lives were right censored at the end day of the experiment. For experiments with antibiotic pulse, the half-life was calculated as the time after the antibiotic pulse when plasmid abundance dropped below the plasmid abundance on day 3 (after the pulse).

**Quantifying plasmid dynamics under chemical treatment**

*E. coli* MG1655 carrying pSC101 were grown overnight from glycerol stock in 500 μL LB + 50 μg/ml Kan (24 hours at 37 °C with shaking at 700 rpm, 2mm) to make sure the purity of the population. The populations were cultured in LB in 96-well plates with periodical shaking (every 5 minutes, orbital for 5 seconds and 2mm), with 1:200 (1 μL into 200 μL) daily dilution rate. When treated, the cell cultures were supplemented with 8 ng/mL Rifampicin (Sigma Aldrich, R3501), 80 μg/ml Promethazine (Sigma Aldrich, 46682), 16 μg/ml Maprotiline (Sigma Aldrich, M9651), and 320 μM Phenothiazine (Sigma Aldrich, 88580). Drip plating was performed to determine the plasmid abundance.

**Generation of barcoded strains**

The Keio strains were generously donated by Dr. Meta Kuehn at Duke University. Barcoded plasmids were generated following the protocols described in previous stuides[3]. Briefly, DNA barcode sequences with random base pairs for barcode sequences were synthesized by an external source and introduced into a linearized plasmid backbone by Gibson assembly cloning. Barcodes plasmid libraries were subsequently transformed into NEB® 5-alpha Electrocompetent *E. coli* cells (Cat. #: C2989) generating a library of barcoded plasmids which were extracted from cells after overnight expansion by plasmid miniprep. Plasmid libraries were used to transform individual Keio strains by chemical transformation. Each strain was validated by Sanger sequencing. Each barcode contains two random 18-bp sequences surrounded by Illumina sequencing adapters.

### Plasmid dynamics in synthetic *E. coli* communities

**Passaging experiment.** *E. coli* Keio strains with barcoded plasmids were cultured in deep well plates for 24 hours at 37 °C with shaking at 700 rpm and amplitude of 2mm from glycerol in 500 µL LB supplemented with 100 µg/mL Carb (for Comm87 strains, **Table S1**) or 25 µg/mL Cm (for Comm57 strains, **Table S2**). Donor strains were cultured for 24 hours at 37 °C with shaking at 700 rpm and amplitude of 2mm in 500 µL LB with double selection for both the barcode plasmid and the target plasmid (10 µg/mL Trim for R388, 10 µg/mL Tetracycline (Sigma Aldrich, 58346-M) for RP4, 100 µg/mL Carb for pCU1, and 100 µg/mL Strp for R6K). The overnight cultures of the plasmid-free strains were mixed in equal proportion, and the mixed recipient community was further mixed with the overnight culture of the donor strain in 3:1 ratio. By doing so, we constructed 5 communities, Comm87 + R388, Comm57 + R388, Comm57 + RP4, Comm57 + pCU1, and Comm57 + R6K. Note for each of the four Comm57 communities, 57 strains are present, including 1 donor strain, and 56 plasmid-free strains. For a plasmid-containing community, e.g., Comm57+R388 having Keio #1 as donor, the community does not include Keio # 2 – 4 which are the donors for the other three plasmids. With initial plasmid abundance of 25%, communities were cultured in LB with either 100 μg/mL Carb (for Comm87) or 25 μg/ml Cm (for Comm57) to prevent contamination and maintain the barcode plasmids, and diluted 1:100 (1 μL into 500 μL) every 24 hours. The four Comm57 communities were also cultured with daily dilution rate of 1:500 (**Fig. S4 & S5**). The plasmid abundance was quantified through selective plating. Cell cultures on the given sampling days were spined down on a benchtop centrifuge, with supernatant discarded, and saved at -20 °C before next-generation sequencing (NGS) library preparation.

**Library preparation**. To prepare the NGS library, nuclease-free water was added to the pellets and samples were boiled at 98 °C for 10 minutes. The barcodes were amplified through PCR (NEBNext Ultra II Q5 Master Mix, NEB, M0544). The primers for PCR each contain unique 8 bp index for multiplexing the samples for sequencing. Products from PCR were pooled together and run on 2% agarose gel. DNA was extracted and purified from the gel using the Zymoclean Gel DNA Recovery kit (VWR, 77001-122), based on the manufacturer's instructions. The cleaned PCR product was compatible with standard Illumina sequencing platforms. The DNA libraries were denatured, diluted, and mixed with an equal amount of Phi-X spike-in based on standard Illumina protocols for preparation of 16S libraries on the Illumina Miniseq. The final libraries contained 50% of Phi-X (Illumina, FC-110-3001) to ensure library diversity. The libraries were sequenced using 300 cycle paired-end reads on Illumina MiniSeq (Illumina, FC-420-1003 or FC-420-1004). The final library is:

5’-AATGATACGGCGACCACCGAGATCTACACXXXXXXXXAACACTCTTTCCC
TACACGACGCTCTTCCGATCTNNNNNNNNNNNNNNNNNNCCTCAGGGTCACTAGG
NNNNNNNNNNNNNNNNNNAGATCGGAAGAGCACACGTCTGAACTCCAGTCACXXX
XXXXXATCTCGTATGCCGTCTTCTGCTTG-3’

Here Ns represent the barcode sequences, and Xs represent sample index.

**NGS data analysis.** A customized Galaxy workflow was used to demultiplex the samples based on the index sequences from the reads and to count the barcodes of each community. The following tools were used: Trim sequences (Galaxy Version 1.0.2+galaxy0), Barcode Splitter (Galaxy Version 1.0.1), FASTQ joiner (Galaxy Version 2.0.1.1+galaxy0).

**Plasmid dynamics in multi-species communities**

**Passaging experiment.** We utilized a previously reported community ‘SynkC’ consisting of 8 sink isolates from a sink p-trap in an intensive care unit from Duke University Hospital[4]. Based on 16S V3/V4 region, they consist of one *Pseudomonas aeruginosa* strain, three *Bacillus* trains, one *Citrobacter* strain, and three *Enterobacter* strains. The 8 isolates and *E. coli* MG1655 were cultured in deep well plates for 24 hours at 37 °C with shaking at 225 rpm and amplitude of 2mm from glycerol in 1000 µL LB media. *E. coli* MG1655 carrying with one of the three plasmids (pSC101, colE1, and pUC) were cultured overnight from glycerol in LB media supplemented with 50 μg/mL Kan for pSC101 and colE1, or 50 μg/mL Spec for pUC.

Since we aimed to achieve habitat partition in sponges, we performed extreme dilution to enable stochastic localization of bacterial cells of each strain within the sponge's porous structure[5]. The overnight cultures of all 10 strains were diluted to OD of 0.3, and mixed together by volume so that plasmid -free and -carrying MG1655 each take up 25% of the total population, and each of the eight sink isolates take up 6.25%. The mixed communities were diluted and inoculated in sponges immersed in 1 mL LB, reaching a further dilution of $4\times10^6$ from OD = 0.3. To further assess the baseline GFP/OD for each plasmid, the three plasmid-carrying MG1655 strains were diluted $1.6\times10^7$ from OD = 0.3 and inoculated in sponges immersed in 1mL LB supplemented with corresponding antibiotics (50 μg/mL Kan for pSC101 and colE1, or 50 μg/mL Spec for pUC). The communities were culture at 37 °C with shaking at 700 rpm and amplitude of 2 mm. The cellulous sponges (ARCLIBER, TBWS-10006) were cut into pieces of 0.55 × 0.55 × 2 cm manually, put into deep 96-well plates and autoclaved for 45 minutes.

For every 24 hours, we squeezed the sponges using pipette tips and extracted the liquid cultures. As above, the cultures were diluted 4-fold (50 μL of culture into 150 μL LB) to measure OD and corresponding GFP intensity. The cultures were further diluted and inoculated in new sponges with final dilution rate of $2\times10^7$. Between day 1 and 2, instead of LB, LB supplemented with corresponding antibiotics was used (50 µg/mL Kan for pSC101 and colE1, or 50 µg/mL Spec for pUC). The same experiment was performed for communities in the absence of the sponge (**Fig. S7**). The extracted liquid cultures were stored in -80 °C before sequenced for their 16S V3/V4 region. 16S sample preparation, sequencing, and analyses were performed by SeqCenter, LLC.

**Sample preparation and sequencing.** Genome DNA was extracted using ZymoBIOMICS DNA Miniprep kit (Zymo research, #D4300). Samples were prepared using Zymo Research's Quick-16S kit (Zymo research, #D6400) with phased primers targeting the V3/V4 regions of the 16S gene. The specific primer sequences are:

| Region | Forward Sequence | Reverse Sequence |
|---|---|---|
| V3/V4 | 341f<br>CCTACGGGDGGCWGCAG<br>CCTAYGGGGYGCWGCAG | 806r<br>GACTACNVGGGTMTCTAATCC |

Following clean up and normalization, samples were sequenced on a P1 or P2 600cyc NextSeq2000 Flowcell to generate 2×301bp paired end reads. Quality control and adaptor trimming was performed with illumina bcl-convert1 (v4.2.4). Primer-dimer sequences identified as PCR artifacts were filtered from the generated FASTQ files based on the following criteria: read length > 150bp, PolyN strings < 10 sequential Ns; PolyG strings < 150 sequential Gs.

**Sequence analysis.** Sequences were imported to Qiime2[6] for analysis. Primer sequences were removed using Qiime2's cutadapt[7] plugin using the following degenerate primer queries:

| Region | Forward Trim Sequence | Reverse Sequence |
|---|---|---|
| V3/V4 | CCTAYGGGNBGCWGCAG | GACTACNVGGGTMTCTAATCC |

Sequences were then denoised using Qimme2's dada2[8] plugin. Denoised sequences were placed into a feature table detailing which amplicon sequence variants (ASVs) were observed in which samples, and how many times each ASV was observed in each sample. Identified ASVs were taxonomically assessed using the Silva 138 99% full-length sequence database[9] and the VSEARCH[10] utility within Qiime2's feature-classifier plugin. ASVs were then collapsed to their lowest taxonomic units (strain, species, genus, etc.), and their counts were converted to reflect their relative frequency within a sample.

### Supplementary figures

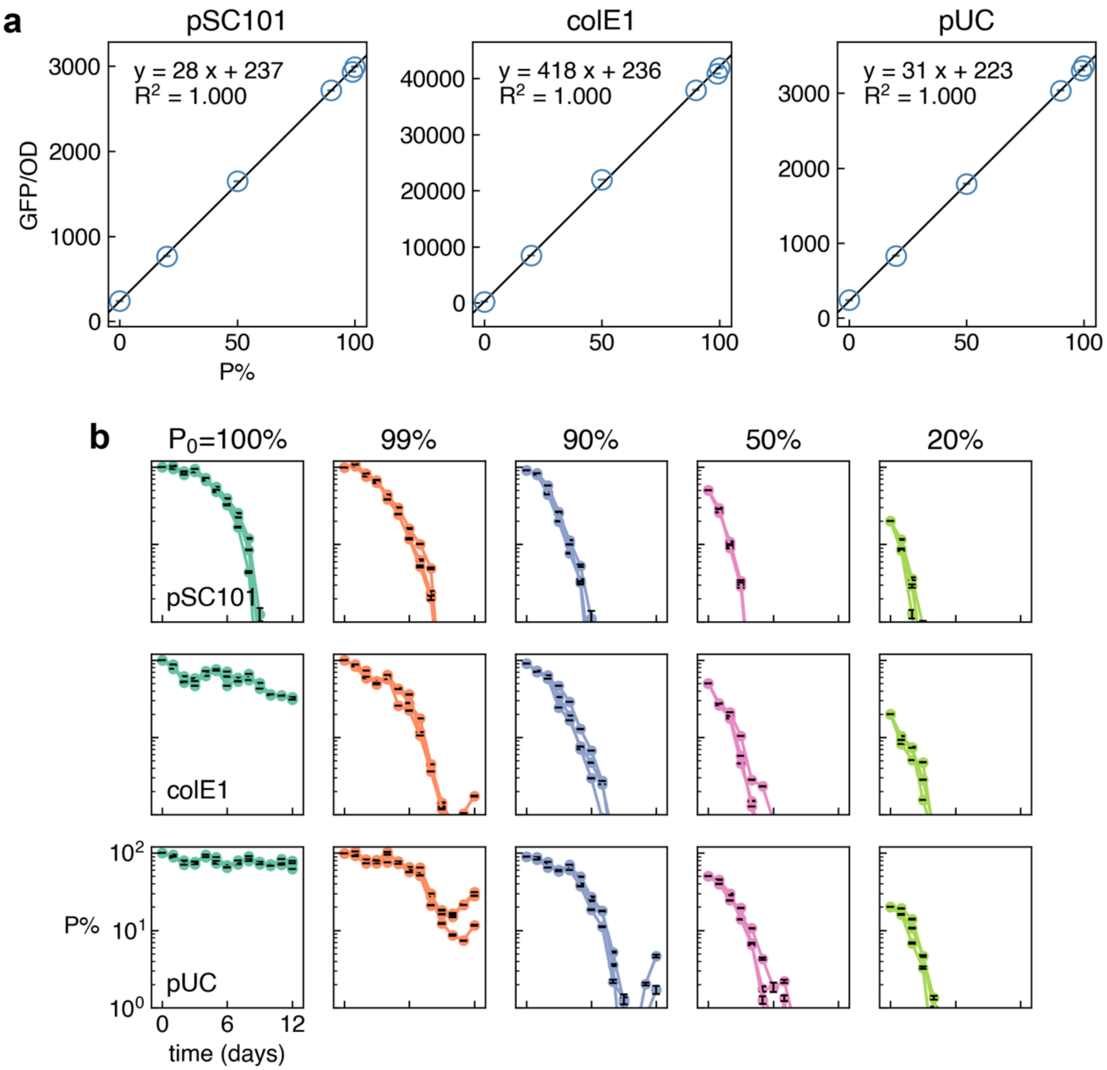


**Figure S1. Measurement of non-mobilizable plasmid dynamics through GFP/OD readout.**

**(a)** Calibration curves for quantifying the abundance of pSC101, colE1, and pUC based on GFP/OD readout. Error bars indicate the uncertainty (SD) in single measurements of GFP/OD. σ-weighted nonlinear fitting was performed to determine the slope and intercept for each calibration curve. pSC101: slope = 27.51 ± 0.03 SE, intercept = 237.16 ± 2.04 SE. colE1: slope = 418.00 ± 0.15 SE, intercept = 235.56± 2.74 SE. pUC:

slope = 31.24 ± 0.03 SE, intercept = 223.49 ± 2.04 SE. 95% confidence intervals (CIs) were also shown but are too narrow to be seen by naked eye. The uncertainties in the estimation of the slope and intercepts were propagated to the calibrated plasmid abundances in **(b)**.

**(b)** Time series of abundance decay for non-mobilizable plasmids in *E. coli* MG1655. Row 1: pSC101, row 2: colE1, row 3: pUC. Colors label different initial plasmid abundance $P_0$%: teal: 100%, orange: 99%, dusty blue: 90%, pink: 50%, and lime green: 20%. Three biological replicates are shown. Error bars indicate propagated measurement uncertainties (per-well SD).

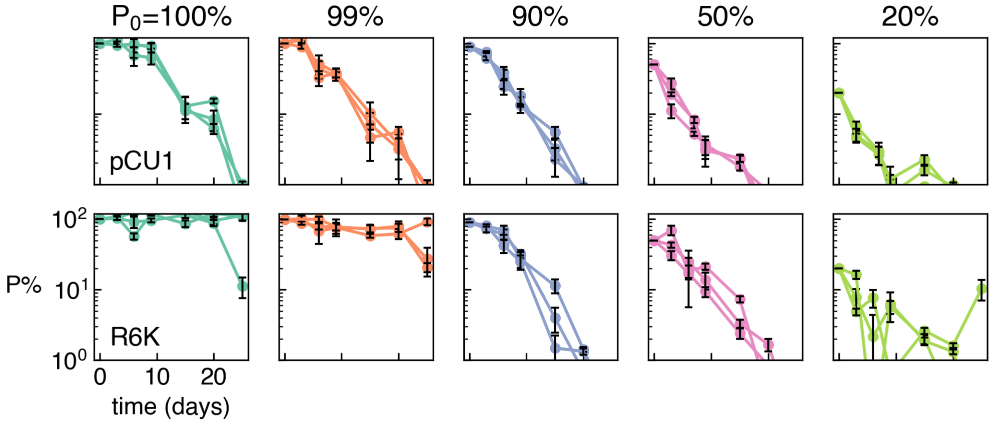


**Figure S2.** Time series of abundance decay for conjugative plasmids in *E. coli* DA28102. Row 1: pCU1, row 2: R6K. Colors label different initial plasmid abundance $P_0$%: teal: 100%, orange: 99%, dusty blue: 90%, pink: 50%, and lime green: 20%. Three biological replicates are shown. Error bars indicate the SE based on technical triplicates of selective plating.

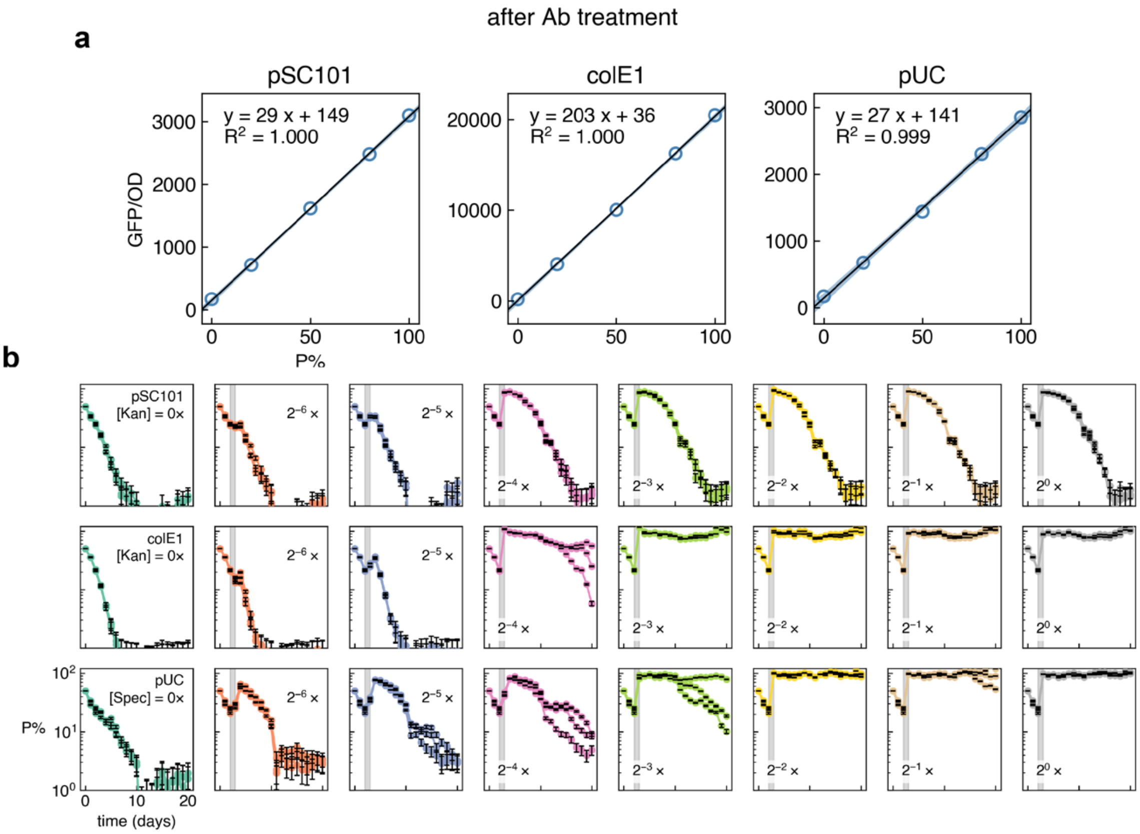


**Figure S3. Non-mobilizable plasmid dynamics following transient antibiotic treatment.**

**(a)** Calibration curves for quantifying the abundance of pSC101, colE1, and pUC based on GFP/OD readout. Error bars indicate the uncertainty (SD) in single measurements of GFP/OD. σ-weighted nonlinear fitting was performed to determine the slope and intercept for each calibration curve. pSC101: slope = 29.33 ± 0.26 SE, intercept = 148.69 ± 16.03 SE. colE1: slope = 203.17 ± 1.72 SE, intercept = 36.14± 107.00 SE. pUC: slope = 26.93 ± 0.41 SE, intercept = 141.33 ± 25.71 SE. 95% confidence intervals (CIs) are shown in grey. The uncertainties in the estimation of the slope and intercepts were propagated to the calibrated plasmid abundances in **(b)**.

**(b)** Time series of non-mobilizable plasmid abundances in *E. coli* MG1655 when treated with transient antibiotic selection. Row 1: pSC101, row 2: colE1, row 3: pUC. Colors label different antibiotic concentrations used between day 2 and 3. Three biological replicates are shown. Error bars indicate propagated measurement uncertainties (per-well SD).

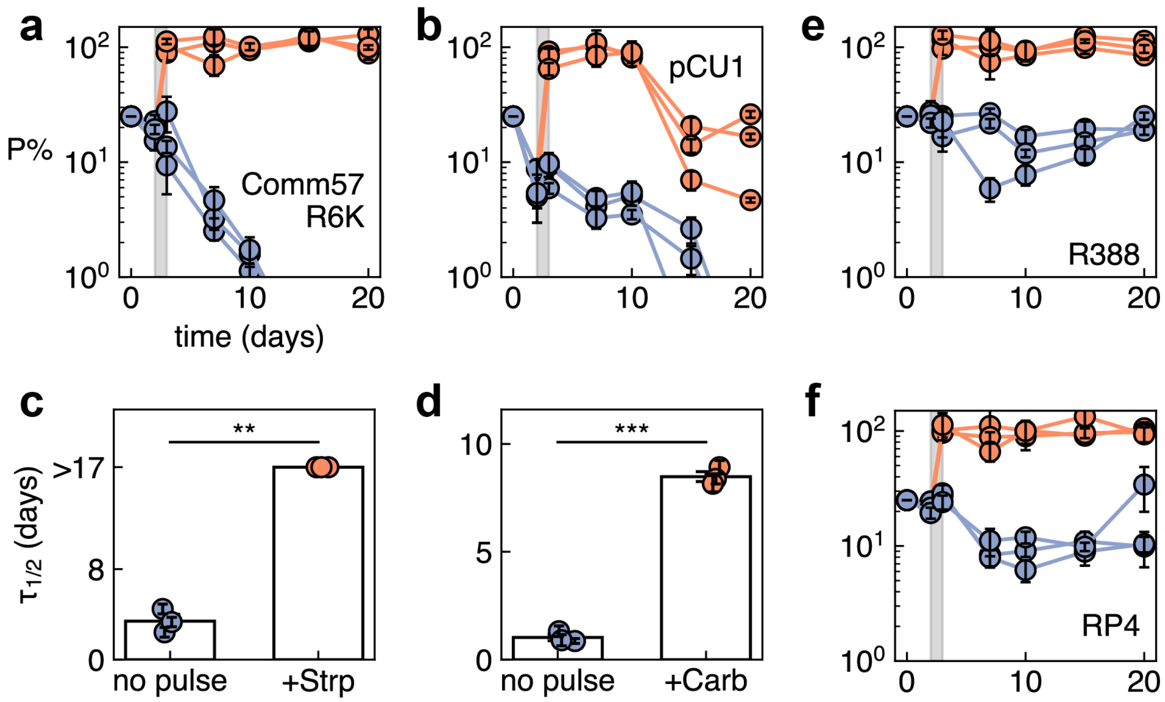


**Figure S4. Antibiotic pulse induces prolonged plasmid carriage in Comm57s passaged at 1:500 daily dilution rate.**

**(a & b)** Plasmid abundance (mean ± SE based on technical triplicates of selective plating) over time for **(a)** R6K and **(b)** pCU1 in Comm57. Purple: communities were passaged daily at 1:500 dilution ratio in LB supplemented with 25µg/mL Cm (Comm57, **a & b**). Orange: communities were treated with additional antibiotic pulse between day 2 and 3 (grey bar) to select for the plasmid-carrying populations (R6K: 100 µg/mL Strp, pCU1: 100 µg/mL Carb). Note for different plasmids, the plasmid donor strains are different while the recipient communities are identical.

**(c & d)** The plasmid half-lives (mean ± SE) were extended by the antibiotic pulses for **(c)** R6K and **(d)** pCU1 in Comm57. Right-censored datapoints ($\tau_{1/2}$ > 17 days) are shown without error bar. Half-lives for R6K in the pulsed group (+Strp) were taken as $\tau_{1/2}$ = 17 days for hypothesis testing. **(c)** Half-lives of R6K after Strp treatment had a higher mean than the LB group: 17.0 ± 0.0 (SD, n = 3) vs 3.4 ± 1.0 (SD, n = 3) days. Welch's t-test:

t(2.00) = 22.83, p=0.002, mean difference = 13.6 [95% CI: 11.0, 16.2], Cohen's d = 18.64. **(d)** Half-lives of pCU1 after Carb treatment had a higher mean than the LB group: 8.5 ± 0.4 (SD, n = 3) vs 1.0 ± 0.3 (SD, n = 3) days. Welch's t-test: t(3.40) = 27.61, p<0.001, mean difference = 7.5 [95% CI: 6.7, 8.3], Cohen's d = 22.55. In the figures, **: p < 0.01, ***: p < 0.001.

**(e & f)** Plasmid abundance (mean ± SE) over time for **(e)** R388 and **(f)** RP4 in Comm57. Antibiotic pulse induced alternative steady state for plasmid persistence. Purple: communities were passaged daily at 1:500 dilution ratio in LB supplemented with 25μg/mL Cm. Orange: communities were treated with additional antibiotic pulse between Day 2 and 3 (grey bar) to select for the plasmid-carrying populations (R388: 10 μg/mL Trim, RP4: 10 μg/mL Tet).

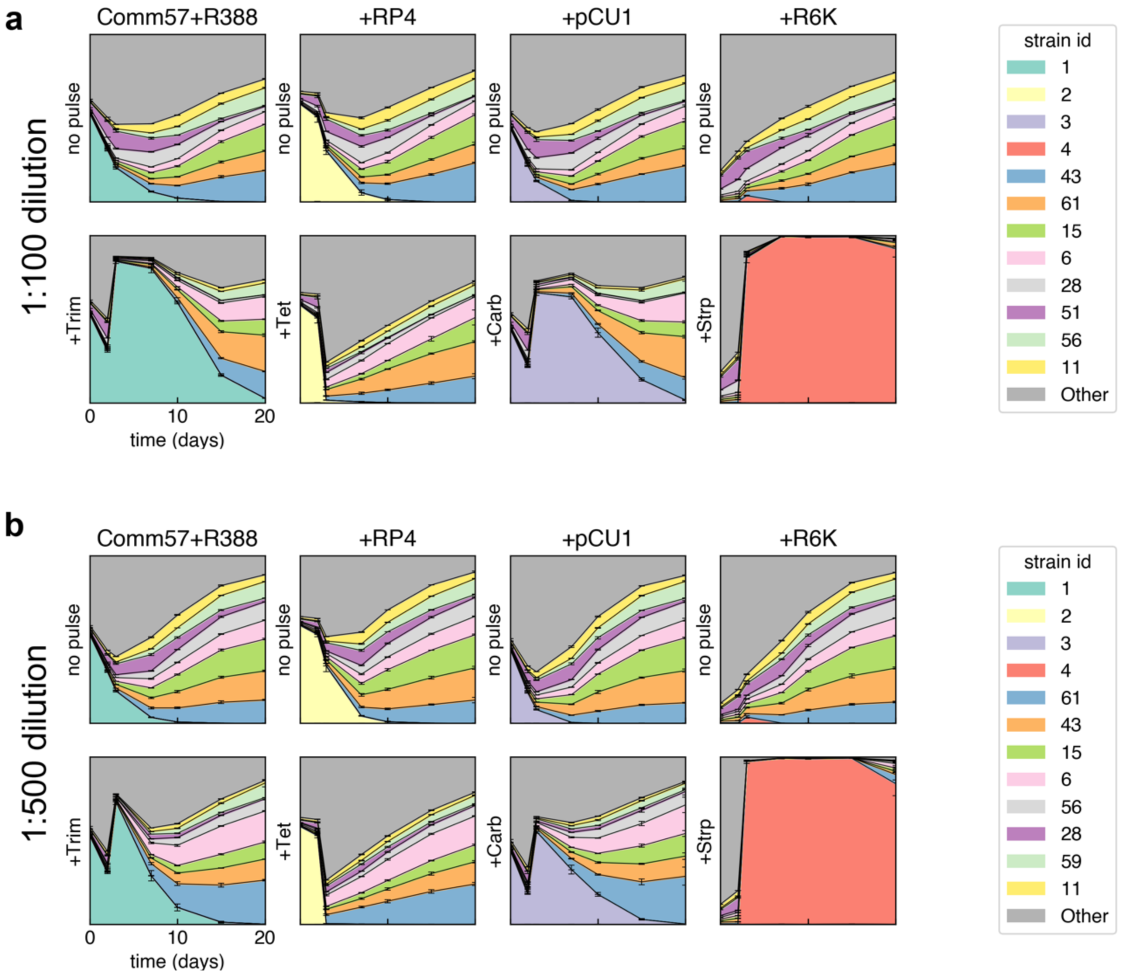


**Figure S5. Strain-level community dynamics of Comm57 with different plasmids.** **(a)**. Strain-level community dynamics (mean ± SE, n = 3) of Comm57 without (top) or with (bottom) 1-day antibiotic pulse, passaged at 1:100 daily dilution rate. The 12 most abundant strains (by accumulative relative abundance across all samples) are color-coded. Note for different plasmids, the plasmid donor strains are different while the recipient communities are identical. Strain 1, 2, 3, 4 corresponds to the donor strain of R388, RP4, pCU1, and R6K.

**(b)**. Strain-level community dynamics (mean ± SE, n = 3) of Comm57 without (top) or with (bottom) 1-day antibiotic pulse, passaged at 1:500 daily dilution rate. Several dominant strains are different from (**a**), so the color codes are slightly different.

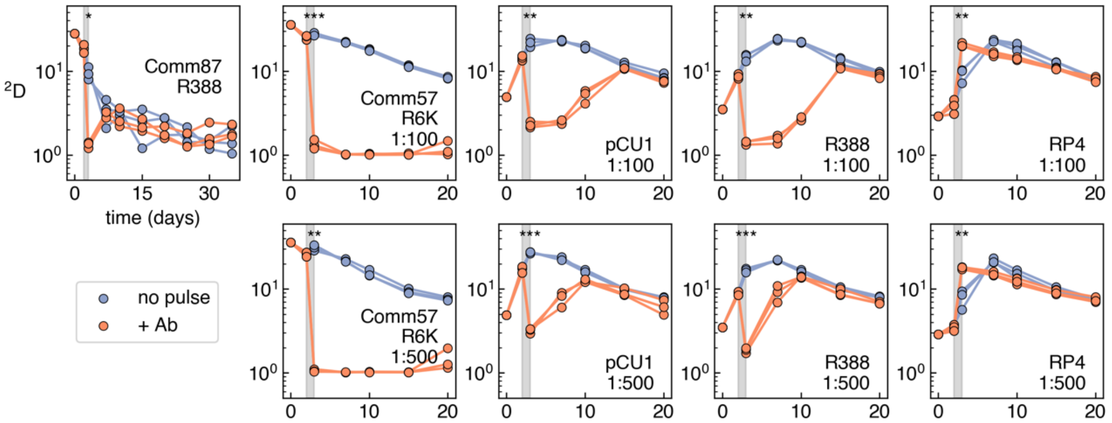


**Figure S6. Community diversity (inverse Simpson index) over time for different synthetic *E. coli* communities.** Statistical significance was determined for the difference in diversity on day 3 (after antibiotic pulse) using Welch's t-test, *: $p < 0.05$, **: $p < 0.01$, ***: $p < 0.001$.

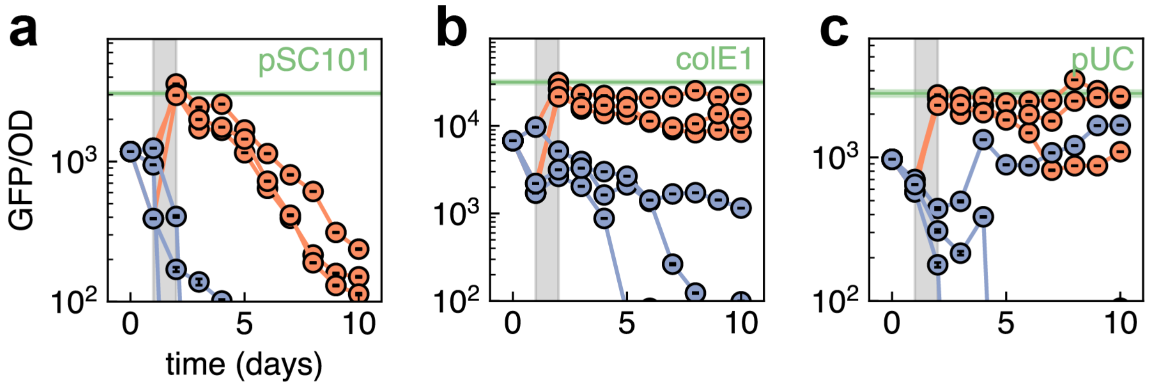


**Figure S7. Ghost effect in multi-species communities of sink isolates.**

Time series of GFP/OD for GFP-encoding plasmids **(a)** pSC101, **(b)** colE1, and **(c)** pUC. Purple: communities were passaged daily at 1:2×$10^7$ dilution ratio in LB. Orange: communities were treated with additional antibiotic pulse between day 1 and 2 (grey bar) to select for the plasmid-carrying populations (pSC101 and colE1: 50 µg/mL Kan, pUC: 50 µg/mL Spec). Green bar indicates the mean ± SE GFP/OD for MG1655 with corresponding plasmid (n = 3). Error bars indicate propagated measurement uncertainties (per-well SD).

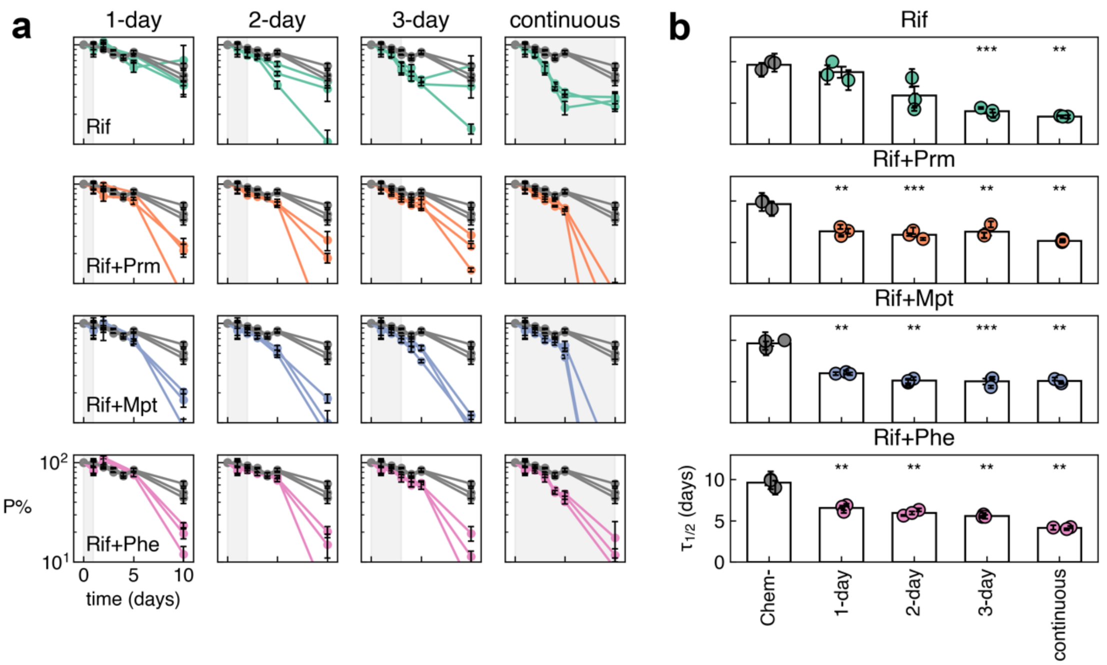


**Figure S8. Plasmid-curing chemicals accelerate plasmid decay.**

**(a)** Time series of non-mobilizable plasmid pSC101 abundances (mean ± SE based on technical triplicates of selective plating) in *E. coli* MG1655 when treated with Rifampicin (teal), Rif + Promethazine (orange), Rif + Maprotiline (dusty blue), Rif + Phenothiazine (pink), and without chemicals (grey). Treatments with different lengths were applied.

**(b)** Plasmid-curing chemicals shortened the half-life of plasmid pSC101. Welch's t-test between the treated groups and the chemical-free control (Chem-), *: $p < 0.05$, **: $p < 0.01$, ***: $p < 0.001$. Right-censored points were taken as $\tau_{1/2}$ = 10 days for hypothesis testing.

**Table S1. Keio strain library and barcodes for Comm87.**

* Donor of plasmid R388.

| ID | Barcode | Knockout gene |
|---|---|---|
| 1* | GGGAAACATGGTTAGAACCCTCAGGGTCACTAGGTAAGAAGCGTACACAAGT | *yncM* |
| 2 | CGGGAAGGTTGGGTTGCTCCTCAGGGTCACTAGGAACATACATGGTACCAGT | *sfcA* |
| 3 | TTATACACCATTCACGAACCTCAGGGTCACTAGGCGAGGGATCGTCATTCCA | *bdm* |
| 4 | CTGGTACTACTAAAAGATCCTCAGGGTCACTAGGCCCAAATGGAAAGTGCGA | *ddpA* |
| 5 | TCATTTGCACTTAAATTTCCTCAGGGTCACTAGGCCATGGCCCACTGGAACG | *yddV* |
| 6 | GAACGGTTCCAGGTTAATCCTCAGGGTCACTAGGCCCATAAACAGATTACCA | *ydeN* |
| 7 | AGAAATTACCACCGTCGTCCTCAGGGTCACTAGGTCCGTGCTACCATGCTGA | *yneE* |
| 8 | TGTTCTACCAGTGTATTACCTCAGGGTCACTAGGCGCCCTCACATACCTATC | *yneI* |
| 9 | CATGCCAGTGGCACTTTGCCTCAGGGTCACTAGGAACGCCCGCTGCCTCTGC | *marR* |
| 10 | CTCCCCGGCAGCTATGGGCCTCAGGGTCACTAGGTTGCTATACACTGAGTGC | *marA* |
| 11 | TCTGATCCCACTCGTCATCCTCAGGGTCACTAGGAATATGCCAGTCTAGCCC | *eamA* |
| 12 | CTATTTTTGTTCGAGCATCCTCAGGGTCACTAGGGTCAGATCAAGCCGTGGC | *ydfO* |
| 13 | GGTCGTTGAGGCGTCCGACCTCAGGGTCACTAGGCCACTACGCAACCCCCAG | *gnsB* |
| 14 | CTTCAATTTAGTTTCGTGCCTCAGGGTCACTAGGCTCGCTCACGAGCGCGCT | *essQ* |
| 15 | ATAGTTCATACGAAGAACCCTCAGGGTCACTAGGCCACCGAGGCGGGCCGAT | |
| 16 | AGAATAAATGAACGCTTTCCTCAGGGTCACTAGGCGAGAGCACGCCGTTCCC | *clcB* |
| 17 | ATAATTTTTAAAGTGGCACCTCAGGGTCACTAGGAGGATGGGAGCCTGCTGG | *ydgJ* |

| | | |
|---|---|---|
| 18 | TATCGCTGTACACATCCTCCTCAGGGTCACTAGGCCCATTCCA CGCCAGGAC | *slyA* |
| 19 | GAGCTATCTACGCCATTTCCTCAGGGTCACTAGGACTCTAAAG CAGCTAAAC | |
| 20 | AGCATTTCGGGAATTTCTCCTCAGGGTCACTAGGTCTCATCAG AACAAACCA | |
| 21 | CCTATTCAAAGCGCAAGTCCTCAGGGTCACTAGGTGTTTACTC ATTATATTT | *ydhO* |
| 22 | AGTTTTGGTATTACTTTGCCTCAGGGTCACTAGGGTGCGGGAA GGCACATAC | *ydhV* |
| 23 | ACTATTACCCGTATTTTGCCTCAGGGTCACTAGGACGTATCAAT ACTATTGG | *sufB* |
| 24 | GGCCCCGGCAGGGCCCCGCCTCAGGGTCACTAGGTACCATAT AACACAATTA | *ydiO* |
| 25 | AAACTTAACTCATCGAATCCTCAGGGTCACTAGGTGCTAAAAG AGCGTTTGT | *ydiQ* |
| 26 | GTAACTTTAGTAGCGGTACCTCAGGGTCACTAGGCCTAGCAAC CCAACATCC | *pfkB* |
| 27 | CTATACGTCATTCTTCATCCTCAGGGTCACTAGGAAGCCAGCT CTGATGTCA | *ydjY* |
| 28 | AGTCAAAGAAATATATCTCCTCAGGGTCACTAGGTGCCTATGG GATCCACGC | *ynjB* |
| 29 | CTAGAAAAACGTGCAATTCCTCAGGGTCACTAGGCTTAATGGT CAATGACTG | *ynjC* |
| 30 | CCGTTTACTGCACACTGTCCTCAGGGTCACTAGGTTGTGCAGC ACCTCGTAC | *ynjE* |
| 31 | TGTTCGGATTCTCACTTACCTCAGGGTCACTAGGTAATTTTTCT ATCTTTTT | *ynjI* |
| 32 | GGAACAGAGCAAGCTAGTCCTCAGGGTCACTAGGCTAACCAT CATACCCCCG | *ydjH* |
| 33 | TGGCATTATGCTTTATTACCTCAGGGTCACTAGGTTACGCTCAT GTCTCAGG | *yeaJ* |
| 34 | AGCGAACGCTATTCTCCCCCTCAGGGTCACTAGGGCTCTAATC ACACAACCT | *yeaP* |
| 35 | TTTCTAAAGGTTCGGTTTCCTCAGGGTCACTAGGACCTACGTA AATAACAGC | *yoaB* |
| 36 | CTCCTCGTGCTATATAAGCCTCAGGGTCACTAGGCACGACTTC ATGGGAAAC | *yoaC* |
| 37 | GCTGCAAGGATTGTATTACCTCAGGGTCACTAGGGGAGTTGAT | |

| | | |
|---|---|---|
| | CAGCGTCCC | |
| 38 | AAAGAAAATAAAAATTTACCTCAGGGTCACTAGGAGCCCAGACGCTCGAGGA | *proQ* |
| 39 | GGAGTTCGTGTTCCATAACCTCAGGGTCACTAGGGTACATAGGTTCTACCAG | *yebU* |
| 40 | CCATTCAGTAAAAAATCTCCTCAGGGTCACTAGGACTTCAGTTCCCGCTCTT | *yebV* |
| 41 | TGGCATGAATACGATTTGCCTCAGGGTCACTAGGAATTAATTCATGGGCGTG | *yebW* |
| 43 | AGAAATGTAATCCTAATACCTCAGGGTCACTAGGAAATCATACCAAATTGGC | *yecD* |
| 44 | GCCGTAAGGTAAATGTGTCCTCAGGGTCACTAGGACTCCGGAAGGACCATCA | *yecN* |
| 45 | GTAACTAACTTCCCTGATCCTCAGGGTCACTAGGCGCAGCACTGAATTTGTA | *yecM* |
| 46 | ATTTTACCTTAAGCGCCCCCTCAGGGTCACTAGGCGACTCTTAACTCCTCCA | *yecT* |
| 47 | GGCTAAAGCAATGTGCGTCCTCAGGGTCACTAGGCTAATCCAGTAGTAATAT | *otsA* |
| 49 | TTCAATACTCTCATTCGCCCTCAGGGTCACTAGGCATTCCAAAGGAAAGCCG | *yodD* |
| 50 | AGTACGTCCTAAGGTGTACCTCAGGGTCACTAGGTGCACAAGACCCATTGCA | *yedS* |
| 51 | TTAGCTTATCTGTCCAATCCTCAGGGTCACTAGGCATCACCTCACGAACCCA | *yodB* |
| 52 | TCAACACATGTGTATGCTCCTCAGGGTCACTAGGGGCCTTCCCAGCCCAGCG | |
| 53 | TGTATAATGTCAATTGGTCCTCAGGGTCACTAGGATTAATACTGACCCTCGG | *dacD* |
| 54 | GAAAGTTACCCTACGGATCCTCAGGGTCACTAGGCCTCTGGCGCAATCTCTT | *nudD* |
| 55 | AGCTGGGTGGGCGGTTTGCCTCAGGGTCACTAGGCTAACTACCCTACATAAC | *yegH* |
| 56 | TAAAGACGACAATTGATGCCTCAGGGTCACTAGGCCACCTTTTGATCTAGTC | *mdtA* |
| 57 | GCCTTTTTCGTCTTTTTGCCTCAGGGTCACTAGGCGGTACCCTCGCCATGCT | *yegP* |
| 58 | TCATTCTTTTCTGTCTGTCCTCAGGGTCACTAGGCCTATGTTCCAGACCAGA | *gatR* |

| | | |
|---|---|---|
| 59 | CTTGTCGGATCTATTACTCCTCAGGGTCACTAGGACCGACCGC CTACACCAC | *fbaB* |
| 60 | TTGTTATAAATAATACCTCCTCAGGGTCACTAGGTTTGCATGTT AATCGGTT | *yegX* |
| 61 | TGCCCCCTAGGCATCAAACCTCAGGGTCACTAGGAATCACAAA CACGCTGAG | *yohN* |
| 63 | GTTTTCCCGTTAAAGTCGCCTCAGGGTCACTAGGTTTTAGTCTT GACACAGC | *yehP* |
| 64 | ACACATAAGGACGTTTATCCTCAGGGTCACTAGGTGTACCTAG GCTAGCTCC | *yehR* |
| 65 | ACCCTCCCGATTTTCTCACCTCAGGGTCACTAGGGATCAAGAA AGCGAGCAC | *yehT* |
| 66 | CGGTGAATATATGCGATGCCTCAGGGTCACTAGGGATAGTACC CCGACTCGA | *yehU* |
| 67 | AACAAATTTCATTTTGTACCTCAGGGTCACTAGGCTCGGTAAG AGTCATCTA | *pbpG* |
| 68 | CGCGCCGGGCCGGCGGGGCCTCAGGGTCACTAGGCCGCGG GCGGAAAATTTA | *yohC* |
| 70 | TTATATCGTTGGTAAGTACCTCAGGGTCACTAGGTTTCGACGT TCCTAACTA | *yeiW* |
| 71 | TTCGGGATGGCACTGGAACCTCAGGGTCACTAGGGCCACTAT CGTTGATTGA | *yeiP* |
| 72 | AGTACTGTATCATACATTCCTCAGGGTCACTAGGCGCACACCG TCCAAAACG | |
| 73 | CTATGCTCAGCCCATGTTCCTCAGGGTCACTAGGTACTGTCAA GGACTCGGC | *yfaZ* |
| 75 | CGTAGGGTGTTTGTTACACCTCAGGGTCACTAGGTCATCTCGC GCACAATTC | *yfbJ* |
| 76 | TTTCGAAATCTTTGCAGACCTCAGGGTCACTAGGTTAGCCTAA GATTCATGG | *yfbT* |
| 77 | TATGTTAGGTTCGATGAGCCTCAGGGTCACTAGGCCAAATCAA ACCCTCAGA | *yfcE* |
| 78 | AGCCAGCGCGGGCGGGGGCCTCAGGGTCACTAGGAAAGACG GACGCCATCCA | *trmC* |
| 79 | GGTTAACGACATGCTGCGCCTCAGGGTCACTAGGGCGCGCTG CATGTCAAGC | *yfcM* |
| 80 | GTTCAGCCAGCACTTTTTCCTCAGGGTCACTAGGCTACTCGCC TAGGCTACA | *yfdL* |
| 81 | AATAGGGGGAGGTCGGGCCCTCAGGGTCACTAGGCCTCAATT | *ypdA* |

| | | |
|---|---|---|
| | TGGACGAAGT | |
| 82 | GGAGGTGAGGGGGAGGGGCCTCAGGGTCACTAGGATCACGGCCCGGTTGGGC | *ypdH* |
| 83 | TTATGTCAAATAGGCCTGCCTCAGGGTCACTAGGACACATACGAAAAACCAA | *yfeA* |
| 84 | AAATTGGTGGTTTGTGCTCCTCAGGGTCACTAGGTTCAAGGGAAACATCATA | |
| 85 | TGATATGATTTCGGGTATCCTCAGGGTCACTAGGACTTACTGAAAGGGGACA | *yfeW* |
| 87 | GTTTCATGCTAACGATGTCCTCAGGGTCACTAGGCCGATTAAAACAGGTACG | *yfgJ* |
| 88 | GGGTTTTCGTGGAGATGGCCTCAGGGTCACTAGGCAATATCAAAATCTGTTC | *sseB* |
| 89 | AGTAACTATATTATCGGCCCTCAGGGTCACTAGGGCTACAAGGAATTTGCAT | *yphG* |
| 90 | TAAGGTAGTTTTAAAAATCCTCAGGGTCACTAGGATACCTCACAAACCCCGA | *yphH* |
| 91 | AGCGTCGAGATGGATGTGCCTCAGGGTCACTAGGCCTGCCAACCGAGCATTC | *yfhK* |
| 95 | ATATGTACCGTTTCCACTCCTCAGGGTCACTAGGTGGTACCCGACTTAACAC | *yfjD* |
| 96 | TTGCCGTATGGATCTAGGCCTCAGGGTCACTAGGGGTCGCCTCACTCTCACT | *yfjO* |

**Table S2. Keio strain library and barcodes for Comm57.**

* Strain 1 – 4 correspond to donors of plasmid R388, RP4, pCU1, and R6K.

| ID | Barcode | Knockout gene |
|---|---|---|
| 1* | GACCTCTCCCACCGCACTCCTCAGGGTCACTAGGGTGGGCTATAGTGAGTAA | *yfjP* |
| 2* | CCCGAATACTTTCGAGCTCCTCAGGGTCACTAGGAGACGATCTCACCGGTAT | *ypjA* |
| 3* | GCTGACTTTAACTTCATTCCTCAGGGTCACTAGGCTAAGATATTTAAACACT | *ypjC* |
| 4* | TTGCATCCGATGACTTGTCCTCAGGGTCACTAGGCAATAACTTCGGTCACTC | *yqaC* |
| 5 | TTAAAATCTAGTATTCCACCTCAGGGTCACTAGGAATAGTGCTCAAACTTGA | *ygaT* |
| 6 | CTACTAATAGGCGAAAATCCTCAGGGTCACTAGGTCGATTGTTACATCATTA | *ygaY* |
| 7 | TTCCCCCCGGCTGCATGGCCTCAGGGTCACTAGGGACAAACCGGCTGTGTCC | *gutQ* |
| 9 | CCTATGGTGCATCGTTTTCCTCAGGGTCACTAGGCACACTATCCCGGCTACA | *ygbF* |
| 11 | GTCAAGTCAATTAATATCCCTCAGGGTCACTAGGAAACTAGGCGGCACATGG | *ygcQ* |
| 12 | ATTCATTTTTGGGGTCGTCCTCAGGGTCACTAGGCACGCAAGACCTAAACCC | *ygcE* |
| 15 | CTAGCAACAACAGCCTATCCTCAGGGTCACTAGGTTACTAAACAACCCGCAA | *ppdB* |
| 16 | ATATGTGGCTTATATACTCCTCAGGGTCACTAGGCAGTCCAGATTAGAAAAC | *yqeH* |
| 17 | TATTCTGCTTTAACTCTACCTCAGGGTCACTAGGGACTGTCGTTGCGCAAGG | *yqeJ* |
| 19 | GAAGTCAATGTTATAAGTCCTCAGGGTCACTAGGAGTGCTCTCTTAAAACAG | *pbl* |
| 20 | TTTCTTAACGAAATCTTACCTCAGGGTCACTAGGAGAAAAGGGCCATAGCAC | *ygeK* |
| 25 | TCTTATGGTAGGCATGTTCCTCAGGGTCACTAGGACCTCATCCCTTAGCGGC | *mltC* |
| 26 | TTCTCTCATAATACACTCCCTCAGGGTCACTAGGGGGGGTCTAGGACCACAC | *tolC* |

| | | |
|---|---|---|
| 27 | TGTCTAGTGACGGAACACCCTCAGGGTCACTAGGTAGCCGAC AAAGTCCTGG | *yhbO* |
| 28 | TTTCCCTACAGTTATGACCCTCAGGGTCACTAGGAAAACTGCT CATCACTGA | *deaD* |
| 30 | CTTGGGGGGGATGGTTGTCCTCAGGGTCACTAGGGTATCACA TTATACTTCC | *yigZ* |
| 31 | TAAACTGGGCGGGAAAAGCCTCAGGGTCACTAGGCCACAATA ACACCCAGAC | *envC* |
| 32 | TATCTGTGTTGGTAGCATCCTCAGGGTCACTAGGGATTCTAAG ATAGGCGCT | *gspD* |
| 33 | TGAGATTGCTTGGTCTCGCCTCAGGGTCACTAGGCAAATGCAT GTACTTTTC | *sgcX* |
| 36 | GCAGTGCGTTTATTCGTGCCTCAGGGTCACTAGGGTAGCCAAT TCCAAGCTT | *hyfI* |
| 43 | GAAGTCAATGTTATAAGTCCTCAGGGTCACTAGGGTTGGCAGC GACGACGAA | *rhsA* |
| 44 | TAGAGGGGTGCGCCTTGACCTCAGGGTCACTAGGTAAAACAG CCACCATAAC | *ybfO* |
| 45 | TTTCGTTCAAATTGGGGTCCTCAGGGTCACTAGGCCACTAGCT CGAAGCAAT | *rhsC* |
| 46 | TTCTGAGCTGATGATACACCTCAGGGTCACTAGGCCCTCACCG GGGCTCCTA | *rhsB* |
| 49 | TTTATTATTCTTCTTGAACCTCAGGGTCACTAGGTGGCACTTTA AATCCGAG | *ybfD* |
| 50 | GTCCGGTGCGGGGGGGGTCCTCAGGGTCACTAGGTATCTGGT GTGATTCACA | *ydcC* |
| 51 | AATCCCTTTTCTGAAAGTCCTCAGGGTCACTAGGCTCAAAACT CCCAGTTGA | *yhhI* |
| 53 | TGATATGTCTATCGGGGGCCTCAGGGTCACTAGGCCATCTCCA ACAAATTCC | *prpB* |
| 54 | AACACCGTGAGTCTTTAACCTCAGGGTCACTAGGGCGTCCGG CAAAAGTCTA | *ydjQ* |
| 55 | TGTAGATTAGATTTTTATCCTCAGGGTCACTAGGCACTCCCCG CACATACGA | *yeaN* |
| 56 | AATAATAGCGAATGGCATCCTCAGGGTCACTAGGAGCAGTTTT CCTCGACTC | *yccK* |
| 57 | GGATTCCAAATTAACGTCCCTCAGGGTCACTAGGAGAAGCCAC CTAGTATTG | *ycjD* |
| 58 | ATTGTATATATTAGATTACCTCAGGGTCACTAGGAACTGACATA | *yebS* |

| | | |
|---|---|---|
| | CTCCTCAC | |
| 59 | GTTTCGCTTGTCTTAACTCCTCAGGGTCACTAGGGTATCTGAG ACGATGCAC | *yphB* |
| 61 | CAAAACTGAGCAATGCTGCCTCAGGGTCACTAGGGTTACTTCT CAACTAGCT | *yagX* |
| 64 | TCTCTTACATTTTTATTCCCTCAGGGTCACTAGGGCGGCTTACC CCGAACAT | *yedS* |
| 65 | ACTCCATAATCCTAATCCCCTCAGGGTCACTAGGGCAAGCGGA CTATTTAAA | *ygcK* |
| 67 | GGACACTCAAGGCCCAGTCCTCAGGGTCACTAGGTGCCTGGC AATGACATTC | *modA* |
| 68 | GATTAGGTATTATCCAGTCCTCAGGGTCACTAGGTCGACTCTT CTACTGATA | *malT* |
| 69 | AAAGACCTAATCCCTACACCTCAGGGTCACTAGGACAAAAACT GTTAAGGAG | *glpR* |
| 74 | CGCCGGAGATCTACAATACCTCAGGGTCACTAGGTCTAAAGCA CGCGTCAAA | *ubiH* |
| 75 | AATATACGACCTGTACAACCTCAGGGTCACTAGGGTAGAACAC TGGCTTACC | *gudD* |
| 76 | TCAACTGTGGGCGCTATTCCTCAGGGTCACTAGGACAAAAACT GTTAAGGAG | *phnM* |
| 78 | TCCTTTTTTAATGAGCTACCTCAGGGTCACTAGGAATAGCCAAA GCACGTTG | *glyS* |
| 80 | TCCGTCTTTATGTTTTCACCTCAGGGTCACTACCTATTAGCACC TGCTGAAC | *sseA* |
| 81 | CCTCTATTCTCGAGCATACCTCAGGGTCACTAGGCACACCGTG ACACAAACA | *narU* |
| 82 | CCCGTATTTACAGCGCACCCTCAGGGTCACTAGGGCAGACTAT ATTTGCTCT | *yhfZ* |
| 84 | GGTTGCGCTTTATTACGCCCTCAGGGTCACTAGGTGTTTTGAG CCCCTATAT | *thrL* |
| 85 | CACGCTTTCTGTTCATGCCCTCAGGGTCACTAGGTACTCCTCG CAACTAGAA | |
| 86 | TATGTCGAACTGGAAGCCCCTCAGGGTCACTAGGGAAATCAAT TATTAGTCA | *ylcG* |
| 87 | GCAAAGGCAATGATGGATCCTCAGGGTCACTAGGTTTTGTGGA ACCGACCCG | *dsbB* |
| 88 | TATGAGTTTTATTAGCCTCCTCAGGGTCACTAGGCCATGGCAT TTTGAATCC | |

| | | |
|---|---|---|
| 90 | AGAGTCTCAAATTTCCGTCCTCAGGGTCACTAGGCAAACCCGCCCCCGGCTA | *kptA* |
| 93 | TGAGATCGACATGTGGGGCCTCAGGGTCACTAGGTCGTAGGAATGATTATTA | *recN* |
| 94 | TAACGAGATAAGTACGCTCCTCAGGGTCACTAGGCCTAATTATGGTTTTCGG | *mutS* |
| 95 | AGCTCAGGTTTGGACTTTCCTCAGGGTCACTAGGTTAAGCATCTCCCCTCCT | *mutH* |

### Supplementary note

#### Analytical derivation of plasmid half-life in clonal population

To investigate the plasmid dynamics in the clonal population, a previous model describing the conjugative plasmid dynamics was used.[1]

$$\frac{df}{dt} = \alpha\mu\left(1 - \frac{f+p}{c}\right)f - Df + \kappa p - \eta f p$$

$$\frac{dp}{dt} = \mu\left(1 - \frac{f+p}{c}\right)p - Dp - \kappa p + \eta f p$$

One key assumption for the derivation is that the total biomass is approximately constant $s = f + p$. Under this assumption, the ODE equations above describing the dynamics of plasmid -free and -carrying cell population densities can then be transformed into population fractions $F = f/s,\ P = p/s$.

$$\frac{dF}{dt} = \alpha\mu\left(1 - \frac{s}{c}\right)F - DF + \kappa P - \eta s F P$$

$$\frac{dP}{dt} = \mu\left(1 - \frac{s}{c}\right)P - DP - \kappa P + \eta s F P$$

Taking $\delta = (\alpha - 1)\mu\left(1 - \frac{s}{c}\right)$, $\mu_p = \mu\left(1 - \frac{s}{c}\right)$, and $\eta^e = \eta s$, the equations become:

$$\frac{dF}{dt} = (\mu_p + \delta)F - DF + \kappa P - \eta^e F P$$

$$\frac{dP}{dt} = \mu_p P - DP - \kappa P + \eta^e F P$$

Moreover, $F + P = 1$. thus

$$\frac{dF}{dt} + \frac{dP}{dt} = \mu_p + \delta(1 - P) - D = 0.$$

Plug in the equation of plasmid carrying population kinetics:

$$\frac{dP}{dt} = \big(D - \delta(1 - P)\big)P - DP - \kappa P + \eta^e F P$$

$$= \delta P^2 - \delta P - \kappa P + \eta^e (1 - P)P$$

$$= (\delta - \eta^e)P^2 - (\delta - \eta^e + \kappa)P$$

Denote

$$a = \delta + \kappa - \eta^e; b = \delta - \eta^e$$

By substituting $z = 1/P$ and $\frac{dz}{dt} = -\frac{1}{P^2}\frac{dP}{dt}$,

$$\frac{dz}{dt} = -b + az$$

By doing the integration, one will arrive at:

$$P(t) = \frac{aP_0}{(a - bP_0)\exp(at) + bpP_0},$$

where $P_0$ is the plasmid carrying population fraction at time 0. The half-life of plasmid decay is thus

$$\tau_{1/2} = \frac{1}{a}\log\left(\frac{2a - bP_0}{a - bP_0}\right)$$

Now we consider the dynamics of $s$. Under the condition $\frac{ds}{dt} \ll \frac{df}{dt}, \frac{dp}{dt}$, meaning when $f$ and $p$ act as fast variables and can almost cancel each other out, we have:

$$\frac{ds}{dt} = \frac{df}{dt} + \frac{dp}{dt} = (\alpha\mu f + \mu p)\left(1 - \frac{s}{c}\right) - Ds = 0$$

For plasmids which are burdensome, $\alpha > 1$. Denoting $(\alpha\mu f + \mu p) = x(t)(f + p)$, we have

$$\mu \leq x(t) \leq \alpha\mu,$$

$$s = \left(1 - \frac{D}{x}\right)c,$$

$$\delta = (\alpha - 1)\mu\left(1 - \frac{s}{c}\right) = (\alpha - 1)\mu\frac{D}{x} \in [(1 - 1/\alpha)D, (\alpha - 1)D]$$

$$\eta^e = \left(1 - \frac{D}{x}\right)\eta c \in [\eta\left(1 - \frac{D}{\mu}\right)c, \eta\left(1 - \frac{D}{\alpha\mu}\right)c]$$

1 Lopatkin, A. J. *et al.* Persistence and reversal of plasmid-mediated antibiotic resistance. *Nature Communications* **8** (2017). https://doi.org:10.1038/s41467-017-01532-1
2 Cooper, N. S., Brown, M. E. & Caulcott, C. A. A Mathematical Method for Analysing Plasmid Stability in Micro-organisms. *Microbiology* **133**, 1871-1880 (1987). https://doi.org:10.1099/00221287-133-7-1871
3 Wu, F. *et al.* Modulation of microbial community dynamics by spatial partitioning. *Nature Chemical Biology* **18**, 394-402 (2022).
4 Şimşek, E. *et al.* Keystone engineering enables collective range expansion in microbial communities. *bioRxiv*, 2025.2001.2011.632568 (2025). https://doi.org:10.1101/2025.01.11.632568
5 Wu, F. *et al.* Modulation of microbial community dynamics by spatial partitioning. *Nature Chemical Biology* **18**, 394-402 (2022). https://doi.org:10.1038/s41589-021-00961-w
6 Bolyen, E. *et al.* Reproducible, interactive, scalable and extensible microbiome data science using QIIME 2. *Nature Biotechnology* **37**, 852-857 (2019). https://doi.org:10.1038/s41587-019-0209-9
7 Martin, M. Cutadapt removes adapter sequences from high-throughput sequencing reads. *2011* **17**, 3 (2011). https://doi.org:10.14806/ej.17.1.200
8 Callahan, B. J. *et al.* DADA2: High-resolution sample inference from Illumina amplicon data. *Nat Methods* **13**, 581-583 (2016). https://doi.org:10.1038/nmeth.3869
9 Quast, C. *et al.* The SILVA ribosomal RNA gene database project: improved data processing and web-based tools. *Nucleic Acids Res* **41**, D590-596 (2013). https://doi.org:10.1093/nar/gks1219
10 Rognes, T., Flouri, T., Nichols, B., Quince, C. & Mahé, F. VSEARCH: a versatile open source tool for metagenomics. *PeerJ* **4**, e2584 (2016). https://doi.org:10.7717/peerj.2584
